# Neural heterogeneity shapes the temporal structure of human working memory

**DOI:** 10.1101/2025.10.31.684900

**Authors:** Daria Kussovska, Robert Kim, Nuttida Rungratsameetaweemana

## Abstract

Working memory (WM) enables temporary retention of information essential for flexible cognition. Although persistent population activity has long been regarded as a principal mechanism of memory maintenance, continuous single-neuron firing is energetically demanding and difficult to reconcile with the heterogeneous firing properties of cortical neurons. Applying single-trial analyses to a dataset of 902 neurons recorded from 21 neurosurgical patients performing a WM task, we found that maintenance was supported by transient, burst-like episodes of coordinated activity rather than sustained firing. Cross-temporal decoding exhibited localized generalization, and decoding accuracy increased with wider temporal windows, indicating that apparent persistence can emerge from temporally interleaved activity across neurons. We further developed a feature-based, putative cell-type classifier that revealed distinct circuit contributions: pyramidal neurons expressed content in burst-aligned events during maintenance, whereas interneurons were strongly modulated by memory load and behavior. Together, these findings reconcile dynamic and persistent accounts, indicating that human WM can emerge from temporally interleaved, cell-type-specific dynamics that provide a flexible and potentially metabolically efficient substrate for maintaining information over time.

## Introduction

Working memory (WM) provides temporary storage and manipulation of information that sup- ports learning, language comprehension, and goal-directed behavior [1–4]. Individual differences in WM capacity are linked to attention [5, 6], problem solving, executive function, information processing, and inhibitory control [5–11], positioning WM as a core component of human cognition [12]. Much of what we know about WM in humans comes from psychophysics, electroencephalogra- phy (EEG), magnetoencephalography (MEG), and functional magnetic resonance imaging (fMRI) [13–26]. Despite these advances, it remains poorly understood how local circuit dynamics and het- erogeneous neuronal populations with distinct intrinsic and functional properties implement WM computations.

Persistent activity has been proposed as a core mechanism of WM, in which subsets of neurons maintain elevated firing throughout maintenance periods [27–33]. Yet this account is difficult to reconcile with the metabolic cost of sustained spiking. Trial-averaged signals that support persistence can also obscure intermittent, task-aligned dynamics at the single-trial and single- participant levels [34]. In addition, evidence for persistence often gives limited consideration to cell-type specificity and the synaptic interactions that shape cortical dynamics [35–37]. This lack of specificity can imply population homogeneity in WM representations [38–40]. Even if persistent activity emerges within distributed networks, uncovering the underlying circuit dynamics requires single-neuron measurements of local circuits that take neuronal heterogeneity into account [41, 42].

Complementary views posit that WM can be maintained through “activity-silent” mechanisms, in which short-term synaptic plasticity stores latent information without continuous spiking [43–45]. In this framework, brief synaptic changes could alter circuit excitability such that stimulus information is expressed through transient, structured bursts rather than uniform persistent fir- ing. Simulations indicate that burst-like activity can optimize information transmission [46]. In nonhuman primates, brief high-frequency spiking accompanies efficient maintenance during visual WM [47]. Experimental and computational work further suggests that balanced excitatory and inhibitory interactions stabilize WM dynamics, underscoring the role of neuronal heterogeneity [48–53]. In rodents, the propensity of pyramidal neurons to burst supports internally generated memory sequences and temporal encoding [54, 55]. In humans, emerging evidence from intracranial recordings is beginning to reveal circuit-level burst dynamics, contributions of distinct cell classes, and functionally defined subpopulations to cognition [56–59]. Yet it remains unknown to what extent single neurons in the human cortex rely on sustained spiking to maintain information over time and how intrinsic properties of distinct cell classes give rise to heterogeneous, temporally structured WM dynamics.

To address this gap, we analyzed intracranial recordings of 902 neurons from 21 neurosurgical patients performing a WM task [60]. We hypothesized that persistent activity is not a strictly local property but an emergent feature of distributed networks. Within local circuits, WM may be supported by activity-silent mechanisms that maintain a latent memory state, while information is periodically expressed through flexible, structured bursts. We further propose that this functionality depends on heterogeneous firing regimes, such that neurons with distinct intrinsic properties differentially reorganize their activity across task epochs to preserve and transform stimulus representations during WM. We performed single-trial, within-participant analyses and applied dimensionality-reduction techniques to move beyond across-trial averaging. We found that stimulus representations during encoding and maintenance occupy distinct low-dimensional subspaces, consistent with evolving rather than static dynamics. WM maintenance is accompanied by transient, burst-like episodes of coordinated activity aligned to task structure. Using a putative cell-type classification, we further show that pyramidal neurons exhibit relatively stable temporal dynamics across task epochs, whereas interneuron activity is closely linked to load and behavior.

Taken together, our results demonstrate that maintenance of stimulus-specific information in WM does not require sustained spiking and can emerge from interactions among heterogeneous neuronal populations expressed through transient, structured activity. These findings advance mechanistic accounts of WM and underscore the value of single-trial, time-resolved analyses that account for neuronal heterogeneity to uncover the circuit dynamics underlying human cognition.

## Results

### Behavior and single-neuron recordings in a working memory task

To probe circuit dynamics underlying working memory (WM) maintenance, we analyzed single-neuron recordings from patients with epilepsy (*n* = 21) while they performed a Sternberg WM task, using a dataset curated by Kyzar et al. ([60]). On each trial, 1–3 images (load 1–3) were presented during an encoding epoch, followed by a maintenance interval and a probe period. Participants in- dicated whether the probe image had appeared during encoding epoch of the current trial (Fig. 1a). Recordings from the 21 patients yielded 902 well-isolated units (Fig. 1b,i), distributed across the amygdala (AMY; 259 neurons), dorsal anterior cingulate cortex (dACC; 171 neurons), hippocam- pus (HPC; 190 neurons), presupplementary motor area (preSMA; 250 neurons), and ventromedial prefrontal cortex (vmPFC; 32 neurons).

**Fig. 1.**
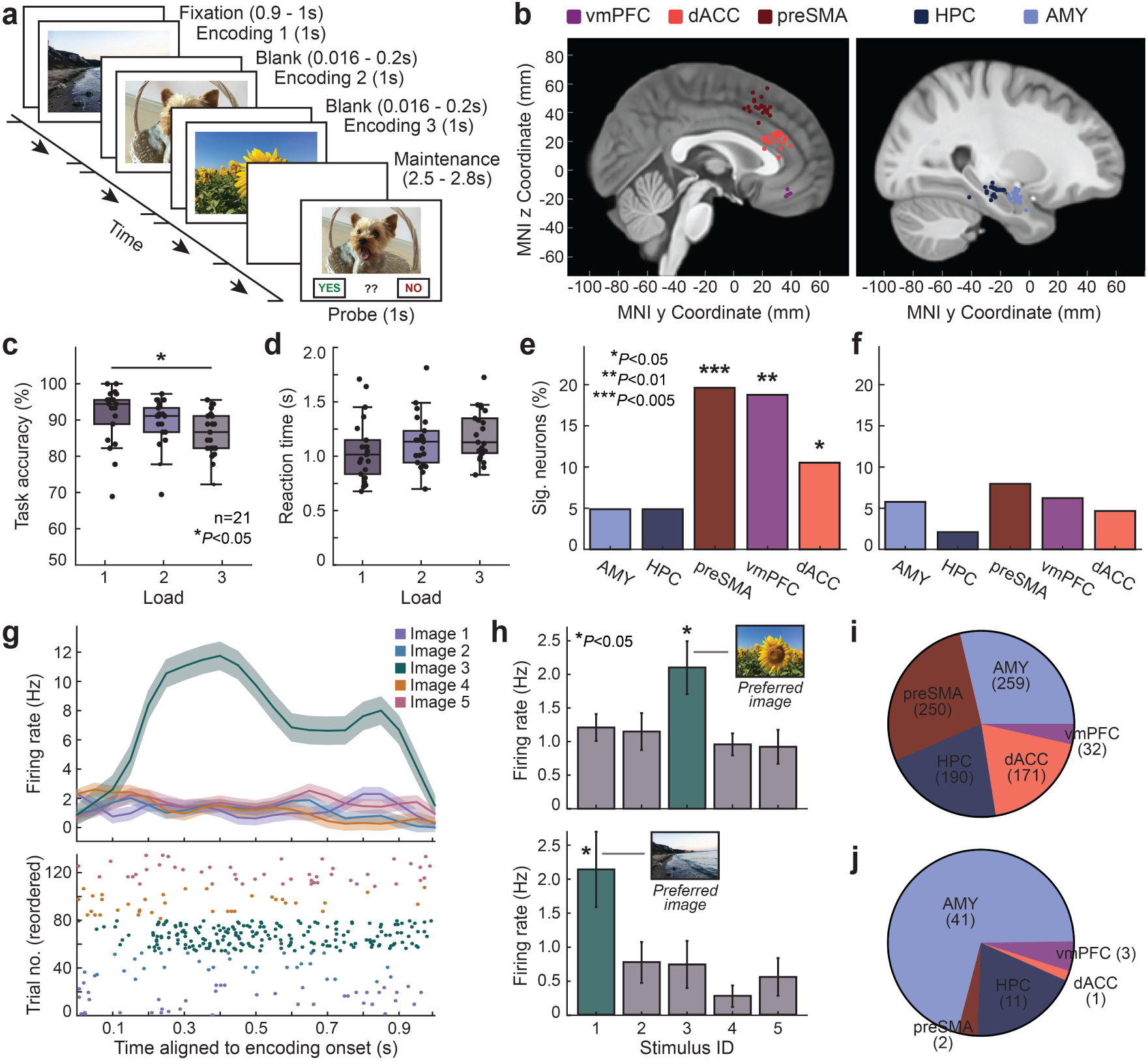
**| Task, recording sites, behavior and concept cell definition a,** Sternberg working memory (WM) task. Each trial begins with encoding of 1–3 images (load 1–3), followed by a 2.5–2.8 s maintenance interval (“Hold”) and a probe requiring an old/new response. If the probe image was presented during encoding on that trial, the correct response is “yes” (old); otherwise “no” (new). Reaction time is measured from probe onset. **b,** Recording sites registered to MNI152 space and overlaid on the CIT168 atlas. Regions assessed are amygdala (AMY), dorsal anterior cingulate cortex (dACC), hippocampus (HPC), presupplementary motor area (preSMA), and ventromedial prefrontal cortex (vmPFC). Microwire bundles across all 41 sessions from 21 patients are shown as individual dots and mapped onto one hemisphere for visualization. **c,** Task accuracy by load across patients. Accuracy is higher for load 1 trials than for load 3 trials (Wilcoxon rank-sum, *P* = 0.027). **d,** Reaction time (RT) for correct trials by load across patients, measured from probe onset. **e,** Proportion of neurons selective for load during the maintenance interval (permutation test, *P*_max_ = 0.026). **f,** Proportion of neurons whose firing rate covaries with RT during maintenance (same statistical procedure as in panel **e**). **g,** Concept-cell criterion. A neuron is classified as a concept cell if it fires significantly more for one stimulus identity during encoding (“preferred stimulus”) than for the other stimulus identities (bootstrap test, *P <* 0.05). Top: peri-stimulus time histogram (PSTH) for encoding (50 ms bins; Gaussian smoothing 50 ms). Bottom: rasters for all trials sorted by stimulus identity. **h,** Two example concept cells: the preferred stimulus (green) elicits higher firing than nonpreferred stimuli (gray) in the encoding window (bootstrap test, *P*_max_ = 0.031). Bars show s.e.m.; the preferred image is shown above each bar. **i,** Recorded neurons by region, aggregated across patients during the WM task (total neurons *n* = 902). **j,** Counts of concept cells by region. Box plots (*c–d*). In each box plot, the center line indicates the median. The box spans the interquartile range, and whiskers extend to 1.5 × interquartile range. \**P <* 0.05; \*\**P <* 0.01; \*\*\**P <* 0.005. All images in this figure are original and created by the authors; no third-party material is included.

Participants performed well on the task (Extended Data Fig. 1a), with high accuracy (Fig. 1c) and fast reaction time (Fig. 1d). Task accuracy was 88.73% ± 1.25% and was higher for load 1 than for load 3 (Wilcoxon rank-sum, *P* = 0.027; Fig. 1c). Reaction time for correct trials was 1.06 ± 0.28 s. Reaction time was faster for probes held in memory than for probes not in memory (permutation test, *P* = 0.001), and correct decisions were faster than incorrect ones (permutation test, *P* = 0.002). Together, these results indicate that participants performed the task as instructed and exhibited typical WM effects [61, 62].

To assess neural population responses across regions during WM maintenance, we analyzed neu- ral pseudo-populations constructed by pooling all recorded neurons across patients (*Methods*). We fit a generalized linear model (GLM) with a separate linear term for each neuron to evaluate sensi- tivity to task variables. During the maintenance epoch, the preSMA, vmPFC, and dACC showed a significant proportion of neurons modulated by memory load (preSMA: 19.6%, vmPFC: 18.75%, dACC: 10.53%; permutation test, *P*_preSMA_ = 0.001, *P*_vmPFC_ = 0.009, and *P*_dACC_ = 0.026; Fig. 1e). We next asked whether neuronal activity scaled with reaction time (RT). Although elevated per- centages of RT-modulated neurons were observed in the amygdala (5.79%), preSMA (8.00%), and vmPFC (6.25%), none were significant after correction for multiple comparisons (permutation tests, *P >* 0.05; Fig. 1f).

### Concept cell definition and criteria

To investigate the neuronal basis of WM, we first identified concept cells. Concept cells were de- fined as neurons selective for stimulus identity, which have been implicated in WM representations in humans [29, 63–65]. For each neuron, we analyzed spiking activity during the first encoding epoch from all trials (load 1–3), counting spikes in a 200–1000 ms window after stimulus onset. In this window, we identified the two images that elicited the highest mean firing rates. To determine stimulus selectivity, we assessed the difference between these two responses using a bootstrap re- sampling test (see *Methods*). Neurons showing a significant difference were categorized as concept cells. For each concept cell, the image eliciting the highest mean firing rate was designated the preferred image.

Concept-cell identification is illustrated (Fig. 1g), and two example concept cells are provided (Fig. 1h). The peristimulus time histogram (PSTH) and raster depict a neuron that responded more strongly to image 3 during encoding, indicating stimulus selectivity (Fig. 1g). This same neuron is classified as a concept cell selective for image 3 (Fig. 1h, top; permutation test, *P* = 0.026). A second concept cell selective for image 1 is also shown (Fig. 1h, bottom; permutation test, *P* = 0.031). Of 902 neurons (Fig. 1i), 58 were identified as concept cells (6.4%; 4 ± 4 across participants). Of these 58 concept cells, 70.69% were in the AMY, 18.96% in the HPC, 1.72% in the dACC, 3.45% in the preSMA, and 5.15% in the vmPFC (Fig. 1j).

### Persistent activity during maintenance emerges with population pooling and coarse temporal binning

After identifying concept cells, we examined how their activity evolves after encoding of the pre- ferred stimulus and whether it takes the form of persistent activity [29, 66]. Consistent with prior work, we predicted that concept cells represent stimulus-specific information throughout the maintenance interval. We also considered that apparent persistence observed when pooling across concept cells may not be present in individual neurons. Because prolonged spiking is metaboli- cally costly, information could be maintained by temporally interleaved activity across different neurons, such that pooled activity within a subpopulation appears sustained. We therefore exam- ined single-neuron activity and population dynamics across task epochs and tested robustness to analysis choices by varying temporal bin width (250–1000 ms in 25-ms steps).

Neuron-by-neuron examination showed that most concept cells did not maintain elevated firing throughout the maintenance epoch (Fig. 2a–b). By contrast, averaging across the concept cell population revealed a clear pattern of persistent activity (*n* = 58; Fig. 2c). This population-level analysis of the Kyzar et al. dataset has been described previously [29, 33]. The effect was robust at loads 1 and 2, and attenuated at load 3 (Extended Data Fig. 1b–c). This dissociation, with heterogeneous single-neuron dynamics alongside population-level persistence, challenges classical WM models that posit continuous persistent firing as the primary mechanism for maintenance [49, 67–69].

**Fig. 2.**
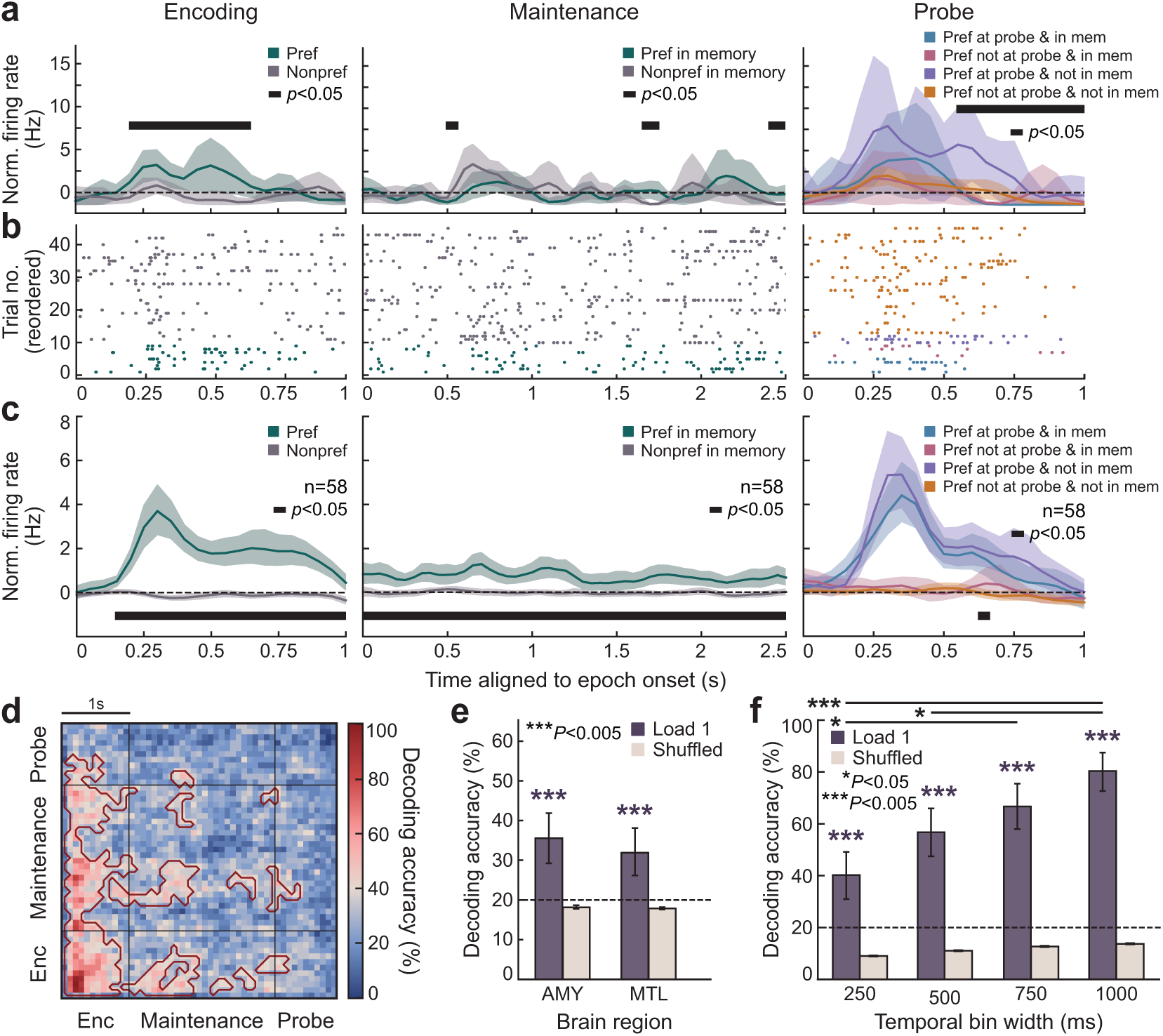
**| Stimulus identity decoding across task epochs. a,** Peristimulus time histograms (PSTHs) from an amygdala concept cell during encoding (left), maintenance (middle), and probe (right). For the encoding and maintenance panels, traces show trial-average, normalized firing rates for load 1 trials when the preferred image was presented versus when the nonpreferred image was presented during the encoding epoch (50 ms bins; Gaussian smoothing 50 ms). Black bars indicate time bins with significant differences between conditions (permutation test, *P <* 0.05). **b,** Raster plots for the same neuron as in **a**, color-coded and sorted by stimulus identity presented at encoding. **c,** Mean normalized firing of all concept cells (*n* = 58) across task epochs for load 1 trials. Shaded bands indicate 95% confidence intervals. Black bars denote time windows with significant differences between preferred and nonpreferred trials (bootstrap test, *P <* 0.05). When averaged, concept-cell activity appears persistent during maintenance. **d,** Cross-temporal decoding for load 1 trials. A linear SVM with leave-one-out cross-validation (LOOCV) was trained and tested on *z*-scored firing rates computed in 250 ms windows stepped every 100 ms. The y-axis denotes training time bins; the x- axis denotes testing time bins. Color indicates accuracy for identifying the encoded stimulus. Red contours mark clusters with accuracy above chance (permutation test, *P <* 0.05). Chance was set to [1/number of stimulus classes] (20%). Cluster sizes expected under the null were estimated from 1,000 label-shuffled iterations and subthreshold clusters were omitted. **e,** Decoding of stimulus identity during the maintenance epoch from concept cells in amygdala (AMY) and medial temporal lobe (MTL), pooled across participants, for load 1 trials. Shuffled-label controls are shown for comparison. The dashed line marks chance (20%). Accuracy exceeded chance in AMY and in MTL (permutation test, *P*_max_ = 0.001). **f,** Decoding accuracy for stimulus identity during maintenance increases with the temporal bin width used for classification. Shown is a linear SVM with LOOCV applied to load 1 trials using bin widths of 250, 500, 750, and 1,000 ms. Shuffled-label controls are shown for comparison. The dashed line marks chance (20%). Asterisks indicate accuracies above chance or differences between conditions (permutation tests, *P*_max_ = 0.001). \**P <* 0.05; \*\*\**P <* 0.005.

To test the stability of population-level sustained activity over time, we performed cross-temporal decoding by training a linear support vector machine (SVM) on pseudo-population activity from concept cells at each time bin and testing it at all other bins, yielding a temporal generalization matrix (see *Methods*). In load 1 trials, decoders trained at specific time points during encoding or early maintenance classified activity at other time points above chance. However, cross-time generalization clusters were sparse and temporally localized, indicating that the stimulus-specific information was not stably maintained across the maintenance epoch but re-emerged at specific moments (Fig. 2d). In load 2 and load 3 trials, cross-time generalization was minimal, suggesting that higher memory load further fragments or destabilizes these representations (Extended Data Fig. 1d–e).

Given our findings suggesting that memory representations over time might not be stable during maintenance, we next quantified the overall amount of stimulus-specific information across the maintenance period, irrespective of its temporal stability. We trained a linear support vector machine (SVM) using a coarse temporal bin width (1 s) on within-participant activity from concept cells during the maintenance epoch, separately for load 1, load 2, and load 3 trials, to classify the identity of the last image presented during encoding (second image for load 2 and third for load 3; see *Methods*). Consistent with prior work, concept cells in AMY and in the medial temporal lobe (MTL; AMY and HPC combined) carried significant stimulus-relevant information during load 1 maintenance (AMY: 35.56% decoding accuracy, *P* = 0.001; MTL: 32.13% decoding accuracy, *P* = 0.001; permutation test; Fig. 2e). The same held for AMY and MTL concept cells at load 2 (AMY: 23.56%, *P* = 0.031; MTL: 28.89%, *P* = 0.002; permutation test; Extended Data Fig. 1f).

Having established that stimulus-related information is present during maintenance, we next asked how varying the temporal window size affects decoding performance. If memory representations are maintained through stable, persistent activity, decoding accuracy should be largely invariant to the bin width used for analysis. However, in line with our cross-temporal decoding (Fig. 2d), we observed that decoding accuracy in the pseudo-population depended strongly on the temporal bin width used to summarize neural activity (Fig. 2f). In load 1 trials, accuracy was above chance at all bin widths (250 ms: 40.00%; 500 ms: 56.67%; 750 ms: 66.67%; 1000 ms: 80.00%; permutation tests against shuffled controls, *P*_max_ = 0.001) and increased nearly twofold as bin width increased. Pairwise comparisons across bin widths confirmed higher accuracies for 750 versus 250 ms (66.67% vs. 40.00%, *P* = 0.026), 1000 versus 250 ms (80.00% vs. 40.00%, *P* = 0.004), and 1000 versus 500 ms (80.00% vs. 56.67%, *P* = 0.032), whereas differences for 500 versus 250 ms (56.67% vs. 40.00%, *P >* 0.05), 1000 versus 750 ms (80.00% vs. 66.67%, *P >* 0.05), and 750 versus 500 ms (66.67% vs. 56.67%, *P >* 0.05) were not significant after correction (permutation tests). Temporal bin width had little effect on decoding accuracy for the second image in load 2 trials. Decoding accuracy in load 2 was also lower than in load 1 (Extended Data Fig. 1g). Together, these results support the view that apparent population-level persistence during maintenance reflects dynamically evolving, temporally interleaved activity across heterogeneous single neurons.

### Encoding and maintenance occupy distinct neural subspaces

To investigate the transient coding suggested by the bin-width and cross-temporal decoding results, we applied demixed principal component analysis (dPCA) to concept-cell activity from load 1 trials. Spike trains were binned in non-overlapping 2-ms intervals, convolved with a Gaussian kernel (*σ* = 60 ms), and *z*-scored relative to the fixation baseline. Encoding and maintenance were analyzed separately, each with its own dPCA fit. For consistency across trials, the maintenance epoch was truncated to 2.5 s. Stimulus separability was quantified as the Euclidean distance between condition-averaged stimulus trajectories in dPCA space, averaged over 100 stratified resampling splits to balance train/test sets (see for further training details and explained variance).

Applying dPCA revealed simple encoding trajectories characterized by a single large excursion in state space, likely reflecting the strong stimulus-evoked response at the onset of the encoding epoch (Fig. 3a). In contrast, maintenance trajectories were markedly more complex, with multiple turns and changes in direction, consistent with evolving rather than static codes (Fig. 3b–c). Stimulus separation, quantified as the mean pairwise Euclidean distance between condition-averaged trajectories in dPCA space, differed significantly across conditions (Kruskal-Wallis, *H* = 457.217, *P* = 1.20 × 10^−97^). Separation in the encoding state space was higher than during maintenance and higher than maintenance activity projected into the encoding state space (Dunn’s post hoc test, *P*_enc vs. maint_ = 2.38 × 10^−38^, *P*_enc vs. projected maint_ = 1.25 × 10^−29^), evident in the large between- epoch differences in distance distributions (see Extended Data Fig. 2i). These results indicate that the maintenance state space differs from the encoding state space, consistent with a represen- tational change between epochs. Projection choice had little effect on maintenance separability: pairwise Euclidean distances among stimulus identities during maintenance were indistinguishable whether activity was projected into the maintenance state space or into the encoding-defined space (Dunn’s post hoc, *P >* 0.05; Fig. 3d). Thus, information about stimulus identity was maintained independent of our chosen projection.

**Fig. 3.**
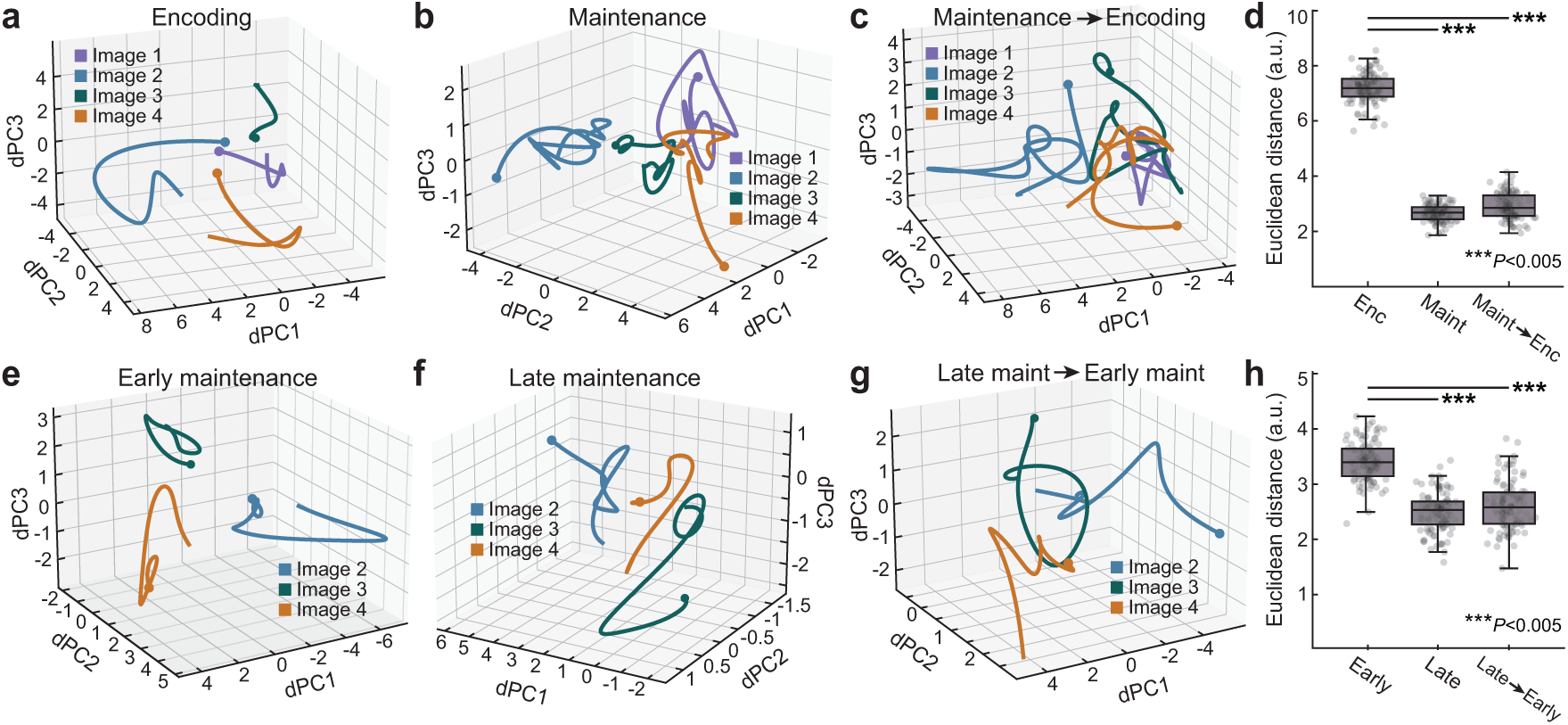
**| Evolving neural population codes across WM epochs. a,** Demixed principal component analysis (dPCA) trained on encoding data and evaluated on held-out encoding test trials. The first three demixed components (dPC1–dPC3) separate the five stimulus identities. Dots mark encoding onset. Colors denote different images (four of the five shown for clarity). **b,** Neural trajectories from held-out maintenance test trials, projected into the maintenance state space, show clear separation by stimulus identity. Dots mark maintenance onset. **c,** The same neural trajectories from maintenance test trials projected into the encoding state space exhibit a different trajectory structure, indicating a representational change between encoding and maintenance. Dots mark the onset of maintenance for trajectories projected onto the encoding state space. **d,** Pairwise Euclidean distances between averaged population responses for each stimulus identity, computed for encoding test data in the encoding state space, maintenance test data in the maintenance state space, and maintenance test data projected into the encoding state space. Separation in the encoding state space is higher than during maintenance and higher than maintenance activity projected into the encoding state space (permutation and pairwise tests, *P*_max_ = 1.25 × 10*^−^*^29^; see *Methods*). **e,** Separation of stimulus specific activity during the first half of the maintenance epoch in held out test data, visualized in the corresponding dPCA state space. Dots mark maintenance onset. Colors denote different images (three of the five shown for clarity). **f,** Separation of stimulus specific activity during the second half of the maintenance epoch in held out test data, visualized in the corresponding dPCA state space. Dots mark the onset of the second half of maintenance. **g,** Projection of activity from the second half of maintenance into the state space defined by the first half, illustrating the evolution of the representation over time. Dots mark the onset of the second half of maintenance for trajectories projected onto the state space defined by the first half of maintenance. **h,** Pairwise Euclidean distances between averaged population responses for each stimulus identity indicate that the early maintenance state space provides greater stimulus separation than late maintenance or late maintenance projected into early maintenance, highlighting temporal changes in the memory representation (permutation tests, *P*_max_ = 2.54 × 10*^−^*^9^; multiple comparisons controlled as described in *Methods*). Box plots: center lines, median; bottom and top edges, lower and upper quartiles; whiskers, 1.5 × interquartile range. \*\*\**P <* 0.005.

This stability, however, was not indicative of a static neural code. When the maintenance period was split into first (early) and second (late) halves (Fig. 3e–f), stimulus separability was significantly higher in the early half than in the late half, and higher than late-maintenance activity projected into the early-maintenance state space (Kruskal-Wallis, *H* = 417.58, *P* = 4.43 × 10^−89^, Dunn’s post hoc, *P*_early vs. late_ = 9.70 × 10^−12^, *P*_early vs. late→early_ = 2.54 × 10^−9^; Fig. 3g–h). The decrease in representational distance over time argues against a stable, persistent code and instead supports a dynamic maintenance mechanism in which stimulus representations are strongest shortly after encoding and gradually reorganize as maintenance unfolds. These findings suggest that apparent population-level stability arises from temporally interleaved activity patterns that continuously reshape the representational landscape during WM maintenance.

### Bursting dynamics reorganize spiking activity between encoding and maintenance

To characterize the evolving code, we quantified population bursts following recent primate and rodent studies [47, 54, 55]. A burst was defined as a transient increase in population activity within a 70-ms bin exceeding a 90th-percentile prominence threshold. To avoid averaging across trials and participants, spikes were concatenated per patient, binned and Gaussian-smoothed (*σ* = 40 ms), and burst candidates within 70 ms of a previous event were suppressed to prevent fragmentation of single events [59]. Parameters were selected by a small grid search (Extended Data Fig. 3a) and are consistent with reports that bursts typically span 50–100 ms [70]. Significance was assessed against a Poisson null model (100 surrogate iterations; see *Methods*) to determine whether observed bursts reflected meaningful neural dynamics rather than random fluctuations (Fig. 4a). Analyses were performed separately for encoding and maintenance to test whether temporal firing patterns differ across epochs. To examine how intrinsic firing properties shape these dynamics, we repeated all analyses in neuron groups stratified by mean autocorrelogram (ACG) amplitude (low vs. high) and ACG decay time constant (*τ*_decay_; short vs. long; Fig. 4b). We used a balanced design in which every participant contributed the same number of neurons for each analysis condition.

**Fig. 4.**
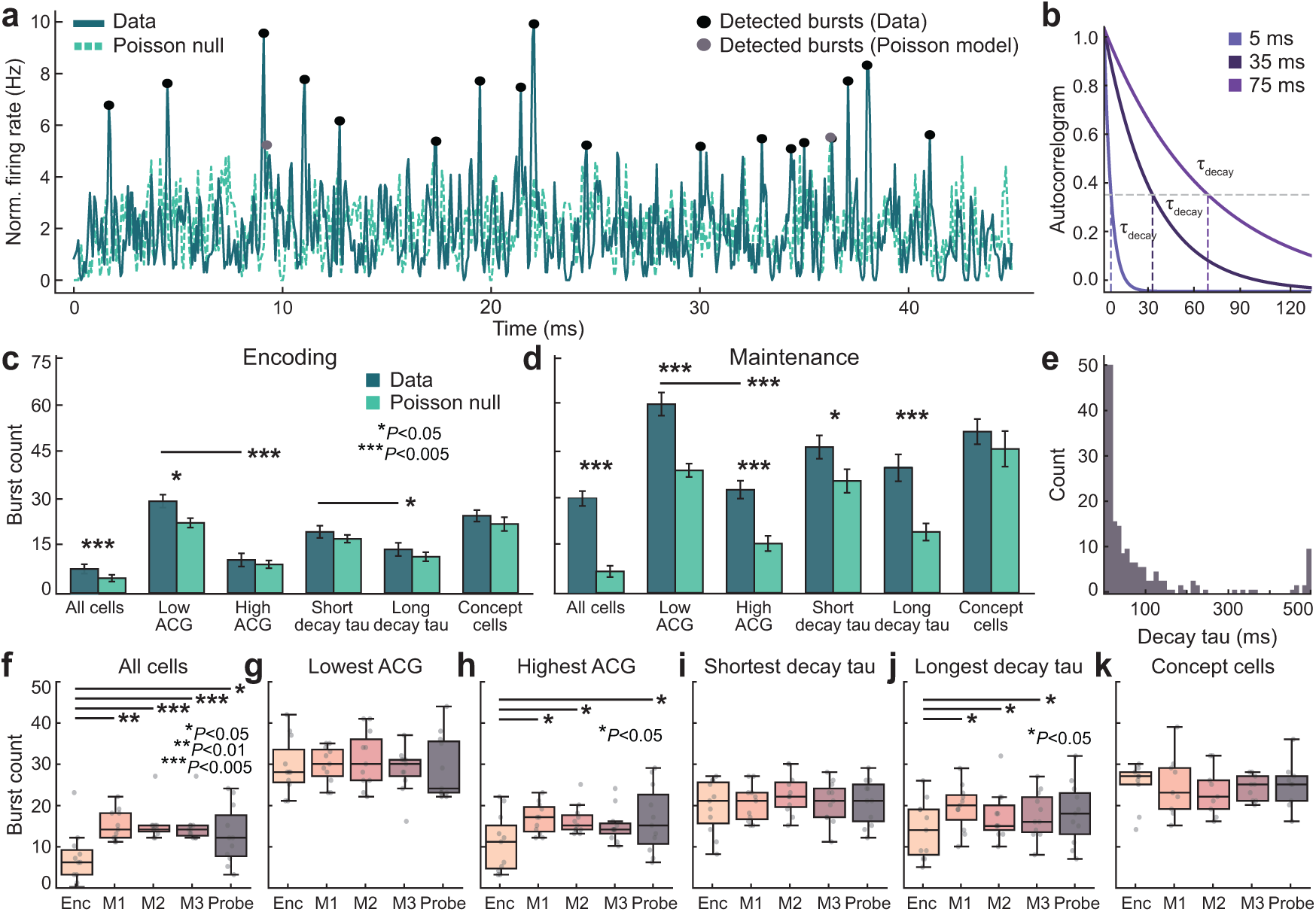
**| Bursting dynamics reorganize from encoding to maintenance. a,** Example burst detection in real spike trains compared with simulated Poisson spike trains matched in rate. **b,** Exponential fits to spike train autocorrelograms (ACGs) illustrating neurons with different decay time constants (*τ*_decay_). **c,** Burst counts during encoding (load 1) in real data compared with Poisson predictions across participants. Deviations from Poisson predictions were observed for all cells (Wilcoxon signed-rank, *P* = 0.003) and in the Low ACG population (Wilcoxon signed-rank, *P* = 0.012; see *Methods* for Low ACG definition). **d,** During maintenance, bursting exceeds Poisson predictions in all neuronal groups except concept cells (Wilcoxon signed-rank, *P*_max_ = 0.011), indicating that elevated bursting in maintenance cannot be explained by random fluctuations. **e,** Distribution of ACG decay time constants (*τ*_decay_) across all recorded neurons. *Panels **f**–**k***. Burst rates by neuronal groups for load 1 trials, comparing encoding, maintenance, and probe. Wilcoxon signed-rank tests were used, with multiple comparisons controlled. Neuronal group definitions, burst detection, and rate normalization are described in *Methods*. **f,** All cells combined: burst rate during maintenance is higher than during encoding, consistent with a task dependent temporal reorganization of spiking (*P*_max_ = 0.039). **g,** Low ACG cells: no significant change in bursting from encoding to maintenance (*P >* 0.05), indicating that low ACG neurons do not drive the increase seen at the population level. **h,** High ACG cells: maintenance shows a robust increase in bursting relative to encoding (*P*_max_ = 0.020), suggesting that high ACG neurons contribute disproportionately to the population-level effect. **i,** Cells with a short *τ*_decay_: no significant difference in bursting between encoding and maintenance (*P >* 0.05). **j,** Cells with a long *τ*_decay_: all maintenance epochs show a higher burst rate than encoding (*P*_max_ = 0.033). **k,** Concept cells: no significant change in bursting between encoding and maintenance (*P >* 0.05). Box plots: center lines, median; bottom and top edges, lower and upper quartiles; whiskers, 1.5 × interquartile range. \**P <* 0.05, \*\**P <* 0.01, \*\*\**P <* 0.005.

During encoding, population bursting did not exceed a rate-matched Poisson null (Wilcoxon signed-rank; Fig. 4c) for most neuronal groups. Significant effects were detected for all neurons (*W* = 9.0, *P* = 0.003) and for the group with low mean ACG amplitude (*W* = 0.0, *P* = 0.012). By contrast, the group with high mean ACG amplitude, the group with a short *τ*_decay_, the group with a long *τ*_decay_, and the concept-cell group did not differ from the null (all *P*_s_ *>* 0.05).

During maintenance, bursting exceeded the Poisson null across most groups (Fig. 4d): all neurons pooled (*W* = 0.0, *P* = 0.001), the low mean ACG group (*W* = 0.0, *P* = 0.004), the high mean ACG group (*W* = 0.0, *P* = 0.004), the short ACG *τ*_decay_ group (*W* = 4.0, *P* = 0.011), and the long ACG *τ*_decay_ group (*W* = 0.0, *P* = 0.004) all showed significant differences, whereas the concept-cell group did not (*W* = 10.0, *P >* 0.05). The absence of significant bursting in the population of concept cells both during encoding and maintenance is consistent with their role in stimulus representation. Their activity is stimulus-locked, not synchronized with the broader network-wide burst episodes, yielding an averaged signal that appears more continuous and less phasic. Together, these results indicate that memory-related population activity is temporally structured, with prominent bursting dynamics that are largely absent during encoding.

Our results revealed that neurons displayed divergent profiles in their burst regulation, indicating distinct functional classes. Specifically, the low mean ACG group showed greater bursting than the high mean ACG group in both encoding (*W* = 0.0, *P* = 0.002) and maintenance (*W* = 0.0, *P* = 0.003), whereas the short *τ*_decay_ group exceeded the long *τ*_decay_ group only during encoding (*W* = 3.5, *P* = 0.015). These findings highlight pronounced functional heterogeneity. Neurons with low mean ACG amplitude show stronger bursting across task epochs, whereas timescale-related differences indexed by *τ*_decay_ are more strongly modulated by task epoch. Thus, ACG amplitude appears to capture a stable intrinsic property of neuronal subpopulations, while *τ*_decay_ primarily reflects task-dependent reorganization of temporal firing dynamics (Fig. 4e).

Motivated by the evolving trajectories between early and late-maintenance identified during dPCA, as well as the population reorganization during maintenance observed from our bursting analysis, we divided the maintenance period into three consecutive 1 s windows (M1–M3) and quantified burst rates over time (Fig. 4f–k). At the population level, burst rates increased from encoding to each maintenance window (load 1; Wilcoxon signed-rank, Enc → M1: *W* = 1.0, *P* = 0.007; Enc → M2: *W* = 0.0, *P* = 0.005; Enc → M3: *W* = 0.0, *P* = 0.005), as well as to probe (load 1; Wilcoxon signed-rank, Enc → Probe: *W* = 4.0, *P* = 0.039). We then asked which neuronal groups drove these changes. Neurons with high mean ACG amplitude increased from encoding to maintenance and to the probe period (Fig. 4h; Enc → M1: *W* = 1.0, *P* = 0.020; Enc → M2: *W* = 3.5, *P* = 0.020; Enc → Probe: *W* = 1.0, *P* = 0.020), and and neurons with a long *τ*_decay_ also increased from encoding to maintenance (Fig. 4j; Enc → M1: *W* = 2.5, *P* = 0.033; Enc → M2: *W* = 2.0, *P* = 0.033; Enc → M3: *W* = 3.0, *P* = 0.033). Other groups showed no change across epochs (Fig. 4g,i,k; Wilcoxon signed-rank, *P >* 0.05). Similar patterns held for load 2 and load 3 (Extended Data Fig. 2b–m), indicating a robust, population-level restructuring driven by specific neuronal subpopulations. Together, these results suggest that WM maintenance relies on dynamic, selectively expressed population bursts rather than uniform persistent firing. Population burst rate increased from encoding into maintenance and across successive maintenance windows. These increases were preferentially driven by neurons with high mean ACG amplitude (Fig. 4h) and long intrinsic timescales (Fig. 4j), yielding a flexible, temporally structured substrate that challenges classical sustained-firing accounts.

### Feature-based classification of putative cell types

Motivated by our bursting results, which showed that neurons with distinct intrinsic properties express different temporal profiles across task epochs, we asked how putative cell types contribute to WM maintenance. Because ground-truth labels are unavailable in human single-neuron recordings, we developed a feature-based, unsupervised classification approach to infer putative identities. We focused on autocorrelogram (ACG) features that capture intrinsic bursting dynamics, providing a functionally grounded alternative to conventional waveform-based criteria (e.g., spike width, asymmetry, amplitude). This allowed us to test directly how single-neuron temporal patterns contribute to population-level bursts. For clustering, we used spectral clustering rather than k- means or Gaussian mixture models, given its robustness to noise and ability to capture nonlinear structure typical of human electrophysiology. This framework offers a novel way to identify putative, functionally distinct neuronal populations that may differentially support WM maintenance.

Prior studies in rodents and non-human primates report characteristic differences in ACG struc- ture between pyramidal neurons and interneurons. Pyramidal neurons typically exhibit a prominent short-lag peak with rapid exponential decay, consistent with intrinsic bursting, whereas interneurons display flatter ACGs and slower decay, consistent with more regular spiking ([71–73]; Fig. 5a–b). For each neuron, we fit Cell Explorer’s triple-exponential model [73] to its spike train and retained well-fit units (*R*^2^ *>* 0.3), yielding 294 neurons for analysis. We extracted three biologically moti- vated features for clustering: firing rate, normalized mean ACG amplitude, and the ACG rise time constant (*τ*_rise_). Given the sparsity of human single-neuron data, we used spectral clustering, which partitioned the population into two clusters (Fig. 5c).

**Fig. 5.**
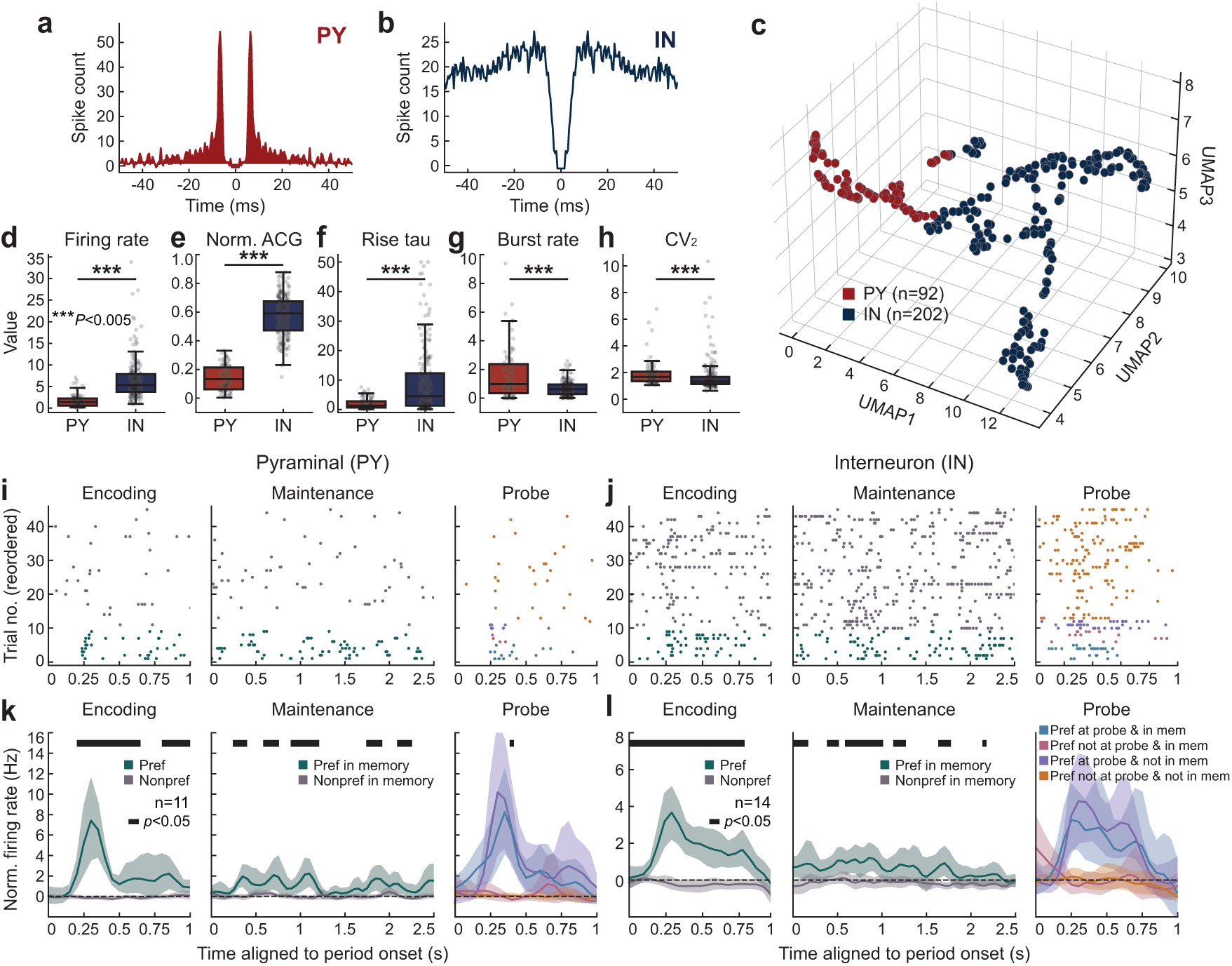
**| Putative pyramidal and interneuron populations show distinct responses during WM. a,** Autocorrelogram (ACG) of an example putative pyramidal neuron with pronounced early peaks reflecting bursting characteristics and a fast exponential decay. **b,** ACG of an example putative interneuron with no pronounced peaks, reflecting regular spiking and slower exponential decay. **c,** Spectral clustering using firing rate, mean ACG, and ACG rise time constant (*τ*_rise_), identified two groups. A UMAP projection for visualization shows points colored by the spectral clustering assignments, corresponding to putative pyramidal (PY, red) and interneuron (IN, blue) populations (PY: *n* = 92; IN: *n* = 202; see *Methods*). **d,** Firing rate by putative cell type. Interneurons had higher firing rates than pyramidal neurons (Wilcoxon rank-sum; *P* = 1.42 × 10*^−^*^32^). **e,** Mean normalized ACG by putative cell type. Interneurons had higher mean ACG than pyramidal neurons (Wilcoxon rank-sum, *P* = 2.16 × 10*^−^*^41^). **f,** ACG rise time constant (*τ*_rise_) by putative cell type. Interneurons had higher *τ*_rise_ than pyramidal neurons (Wilcoxon rank-sum, *P* = 2.58 × 10*^−^*^9^). **g,** Burst rate by putative cell type. Pyramidal neurons had higher burst rate than interneurons (Wilcoxon rank-sum, *P* = 6.35 × 10*^−^*^6^). **h,** Local coefficient of variation (CV_2_) by putative cell type. Pyramidal neurons had higher CV_2_ than interneurons (Wilcoxon rank-sum, *P* = 4.84 × 10*^−^*^4^). **i,** Trial-wise rasters for an example pyramidal concept cell across encoding, maintenance, and probe epochs in load 1 trials, split into preferred and nonpreferred conditions. **j,** Trial-wise rasters for an example interneuron concept cell across encoding, maintenance, and probe epochs in load 1 trials, split into preferred and nonpreferred conditions. **k,** Population PSTH for pyramidal concept cells (*n* = 11) across encoding, maintenance, and probe epochs (50 ms bins; Gaussian smoothing 50 ms; shaded bands show s.e.m.). Horizontal black bars mark time bins with significant differences between plotted conditions (bootstrap test, *P <* 0.05). **l,** Population PSTH for interneuron concept cells (*n* = 14) across encoding, maintenance, and probe epochs (50 ms bins; Gaussian smoothing 50 ms; shaded bands show s.e.m.). Horizontal black bars mark time bins with significant differences between plotted conditions (bootstrap test, *P <* 0.05). Box plots: center lines, median; bottom and top edges, lower and upper quartiles; whiskers, 1.5 × interquartile range. \*\*\**P <* 0.005

Cluster identities were assigned according to established biophysical signatures. One cluster comprised neurons with higher firing rates (Mann–Whitney *U* = 17294, *P* = 1.42 × 10^−32^; Fig. 5d), greater normalized mean ACG amplitude (*U* = 18522, *P* = 2.16 × 10^−41^; Fig. 5e), and larger *τ*_rise_ (*U* = 13465, *P* = 2.58 × 10^−9^; Fig. 5f). We labeled this cluster as putative interneurons (IN; *n* = 202). The remaining neurons, with lower firing rates, lower normalized mean ACG amplitude, and smaller *τ*_rise_, were labeled as putative pyramidal neurons (PY; *n* = 92).

We validated these assignments using independent spiking statistics. Putative pyramidal neu- rons showed higher burst index (PY = 0.985 vs. IN = 0.622; *U* = 6835, *P* = 6.35 × 10^−6^; Fig. 5g). They also exhibited greater spike-train irregularity (CV_2_; PY = 1.673 vs. IN = 1.319; *U* = 6056, *P* = 4.84 × 10^−4^; Fig. 5h), consistent with more irregular pyramidal firing and more regular in- terneuron spiking reported in prior literature [71, 74, 75]. Applied to a labeled rodent dataset [73], the classifier achieved 97% accuracy and reproduced type-specific ACG metrics (Extended Data Fig. 4a–e). These results indicate that the classifier generalizes robustly and can be applied to human data in downstream analyses.

### Distinct temporal activity patterns during memory maintenance in interneurons and pyramidal neurons

Application of the cell-type classification revealed that putative pyramidal and interneuron units showed divergent temporal response patterns during WM maintenance (Fig. 5i–l). Pyramidal concept-cell populations showed pronounced bursting (Fig. 5i,k), whereas interneuron populations exhibited smoother regular-spiking activity (Fig. 5j,l).

We next asked whether these putative cell types differentially contribute to WM maintenance. We compared temporal structure, information content, and task modulation across cell types, starting with bursting. Across participants, pyramidal populations showed elevated bursting during both encoding and maintenance relative to chance (label-shuffled null; Encoding: *W* = 1.0, *P* = 0.012; Maintenance: *W* = 0.0, *P* = 0.004; Fig. 6a–b). By contrast, interneurons exhibited elevated population bursting during maintenance (*W* = 5.0, *P* = 0.001) but not during encoding (*W* = 41.0, *P >* 0.05), suggesting reorganization of spiking dynamics as activity transitioned from stimulus- driven responses to WM maintenance (Fig. 6a–b).

**Fig. 6.**
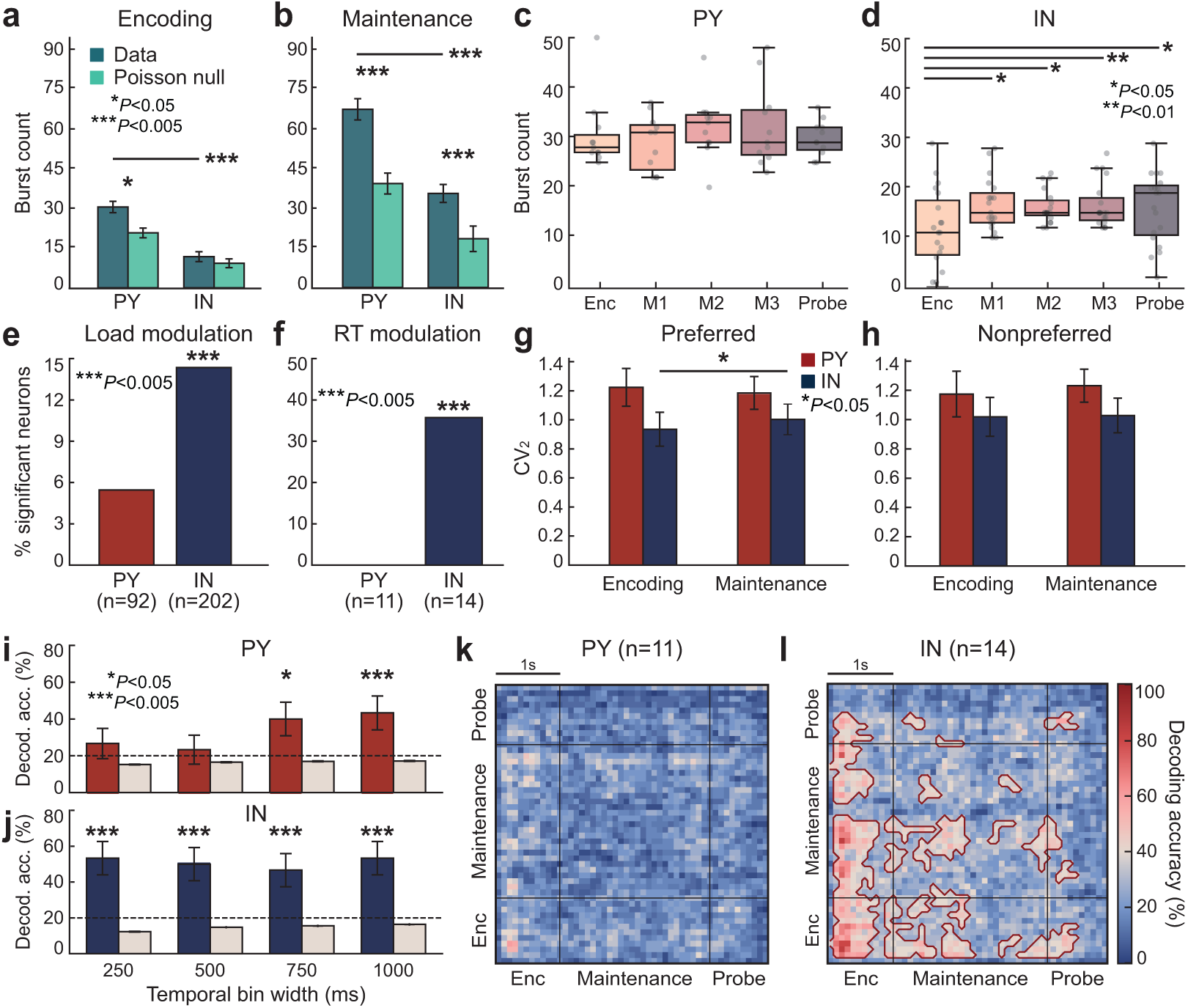
**| Interneurons preferentially exhibit maintenance dynamics and representations. a,** Burst counts during encoding in real spike trains compared with rate-matched Poisson surrogates for pyramidal neurons (PY) and interneurons (IN). PY burst counts exceed Poisson predictions (Wilcoxon signed-rank, *P* = 0.012). **b,** Burst counts during maintenance in real spike trains compared with rate-matched Poisson surrogates for PY and IN. Burst counts in both PY and IN exceed Poisson predictions (Wilcoxon signed- rank, *P*_max_ = 0.004). **c,** Burst counts across task epochs (encoding; maintenance M1, M2, M3; probe) in load 1 trials for pyramidal neurons (PY), where M1, M2, and M3 denote the first, second, and final thirds of the maintenance epoch. The rates remain relatively stable across epochs. **d,** Burst counts across epochs for interneurons (IN). IN exhibit epoch-dependent changes (Wilcoxon signed-rank, *P*_max_ = 0.024). **e,** Proportion of PY and IN with load-modulated firing during maintenance (permutation test, *P* = 0.002). **f,** Proportion of concept-selective PY and IN with reaction time (RT)-modulated firing during maintenance (permutation test, *P*_PY_ *>* 0.05; *P*_IN_ = 0.002). **g,** Local coefficient of variation (*CV*_2_) on preferred trials across encoding and maintenance for concept cells in PY and IN. IN concept cells, but not PY, exhibit a maintenance- related change in *CV*_2_ (Wilcoxon signed-rank, *P* = 0.032). **h,** *CV*_2_ on nonpreferred trials across encoding and maintenance for PY and IN concept cells. No change is observed (Wilcoxon signed-rank, *P >* 0.05). **i,** Decoding of stimulus identity during maintenance from PY concept cells for load 1 trials as a function of analysis bin width (250, 500, 750, and 1,000 ms). Temporal bin width strongly affects PY accuracy. **j,** Decoding accuracy during maintenance from IN concept cells for load 1 trials as a function of temporal bin width. Accuracy remains above chance across widths and is less sensitive to bin width than PY (**i**). *Panels **i**–**j***. Shuffled-label controls are shown for comparison. The dashed line denotes chance (20%). Asterisks indicate accuracies above chance (permutation test, *P*_max_ = 0.030). **k,** Cross-temporal decoding for PY concept cells for load 1 trials. **l,** Cross-temporal decoding for IN concept cells for load 1 trials. *Panels **k**–**l***. Color indicates accuracy for identifying the encoded stimulus. Red contours mark clusters with accuracy above chance (permutation test, *P <* 0.05). Chance was set to [1/number of stimulus classes] (20%). Cluster sizes expected under the null were estimated from 1,000 label-shuffled iterations and subthreshold clusters were omitted. Box plots: center lines, median; bottom and top edges, lower and upper quartiles; whiskers, 1.5 × interquartile range. \**P <* 0.05, \*\**P <* 0.01, \*\*\**P <* 0.005.

Dividing the maintenance period into three consecutive 1 s windows (M1–M3) yielded the same pattern. Pyramidal burst rates remained stable across task epochs and load (Wilcoxon signed-rank, all *P >* 0.05; Fig. 6c, Extended Data Fig. 5a,c), whereas interneuron burst rates changed across task epochs (load 1; Enc → M1: *W* = 17.0, *P* = 0.011; Enc → M2: *W* = 26.0, *P* = 0.024; Enc → M3: *W* = 7.5, *P* = 0.001; Enc → Probe: *W* = 18.5, *P* = 0.003; Fig. 6d). Interneuron burst-rate differences across maintenance epochs increased with load (Extended Data Fig. 5b,d), consistent with greater demands as more items are held in memory.

In addition, interneurons, but not pyramidal neurons, showed modulation of spiking with task parameters. A significant portion of interneurons was modulated by memory load during mainte- nance (14.36%; permutation test, *P* = 0.002; Fig. 6e), including interneuron concept cells (21.42%; *P* = 0.032; Extended Data Fig. 5e). By contrast, pyramidal neurons showed minimal load modu- lation (all PY: 5.43%, *P >* 0.05; concept PY: 0.00%, *P >* 0.05; Fig. 6e; Extended Data Fig. 5e). Interneuron concept cells were also modulated by reaction time (RT; 35.71%; *P* = 0.002), although this effect was not robust across all interneurons (*P >* 0.05; Fig. 6f; Extended Data Fig. 5f). Pyra- midal neurons showed no significant RT modulation in either the full population or the concept-cell subset (permutation tests, all *P >* 0.05; Fig. 6f; Extended Data Fig. 5f).

Our analysis revealed differences in burst dynamics across putative cell types, so we next asked whether these differences are evident at the single-neuron level. Since concept cells carry stronger and less noisy stimulus-specific information, we tested whether their firing patterns differed between preferred and nonpreferred trials. We compared the spike-train local coefficient of variation (CV_2_) in individual pyramidal and interneuron concept cells as a function of whether the preferred image was present in the encoding period of each load 1 trial. Pyramidal concept cells showed stable CV_2_ across preferred and nonpreferred trials during both encoding and maintenance (Wilcoxon signed-rank, all *P >* 0.05). By contrast, interneuron concept cells exhibited higher CV_2_ during maintenance than during encoding on preferred trials (*W* = 13.0, *P* = 0.032), but not on nonpreferred trials (*W* = 50.0, *P >* 0.05; Fig. 6g–h), indicating an epoch-dependent shift in interneuron firing regularity consistent with maintenance-related structure. Firing decay dynamics also differed with stimulus preference. Pyramidal concept cells showed increased decay time constants during encoding and maintenance on preferred trials (Wilcoxon signed-rank, *P*_max_ = 0.029), an effect absent in interneurons (all *P >* 0.05; Extended Data Fig. 5g–h).

We next investigated whether differences in firing patterns between pyramidal neurons and interneurons translated into differences in stimulus decodability during maintenance. For pyramidal neurons, decoding accuracy did not exceed chance with short temporal windows (250 ms: 26.67%, permutation test, *P >* 0.05; 500 ms: 23.33%, *P >* 0.05), likely because these bins were too small to capture sufficient burst events (Fig. 6i). Accuracy improved with larger windows (750 ms: 40.00%, *P* = 0.030; 1000 ms: 43.33%, *P* = 0.030), consistent with larger temporal windows capturing burst-driven activity (Fig. 6i). Cross-temporal decoding showed no above-chance generalization across time even for populations of 11 pyramidal concept cells (Fig. 6k), suggesting that irregular, burst-aligned spiking may underlie maintenance in pyramidal neurons. These findings also highlight a limitation of conventional decoders that do not account for heterogeneous burst timing.

In contrast, interneurons (*n* = 14) decoded above chance across all window sizes (250 ms: 53.33%, *P* = 0.002; 500 ms: 50.00%, *P* = 0.002; 750 ms: 46.67%, *P* = 0.002; 1000 ms: 53.33%, *P* = 0.002; Fig. 6j) and in cross-temporal maps (Fig. 6l), consistent with more regular spiking that yields a reliable preferred-stimulus signal across bins. Interneurons also decoded the second stimulus in load 2 trials above chance across all windows (all *P <* 0.05), an effect not observed in pyramidal concept cells (all *P >* 0.05; Extended Data Fig. 5i–j). Together, these findings indicate a functional dissociation within WM. Pyramidal neurons support selective coding via burst-aligned responses to preferred identities, whereas interneurons exhibit task- and epoch-related modulation consistent with roles in gain control, temporal precision, or network stabilization as load increases.

## Discussion

In this study, we leveraged intracranial single-neuron recordings from 21 human participants performing a working memory (WM) task to examine how heterogeneous neural populations sustain information over time. We found that WM maintenance does not rely on uniform, persistent spiking but instead emerges from dynamic, temporally structured activity across distinct neuronal groups. Apparent population-level persistence arose from temporally interleaved bursts across neurons, while cell-type analyses showed burst-driven responses in putative pyramidal neurons and flexible, load- and behavior-linked modulation in putative interneurons. Together, these findings suggest that human WM is maintained through heterogeneous and cell-type-specific interactions that enable efficient, adaptive coding rather than continuous sustained activity.

A long-standing account of WM proposes that information is maintained through sustained, stimulus-specific firing of single neurons during the maintenance period [27, 29, 32]. Such persistent activity has been reported in nonhuman primates [28, 38] and in humans [29, 33, 66], particularly in prefrontal and medial temporal regions, and is often interpreted as the primary substrate of active maintenance. Our results do not dispute the presence of persistent firing but instead indicate that it is not the sole mechanism supporting memory retention. When activity was averaged across neurons, we observed a population-level signature consistent with persistence. However, single-trial analyses revealed heterogeneous single-neuron dynamics rather than continuous firing in individual neurons. The strong dependence of decoding accuracy on temporal bin width (Fig. 2f, Fig. 6i) further supports a model in which information is expressed through intermittent, burst-like events that, when pooled across neurons and time, yield an apparently sustained representation. These findings reconcile classical models of persistent activity with more recent evidence for dynamic coding, suggesting that population-level persistence reflects structured temporal coordination among diverse neuronal subpopulations rather than uniform tonic firing.

Our findings indicate that the stability of WM does not necessarily depend on continuous firing but rather on coordinated temporal structure within neural populations. Consistent with this view, Panichello *et al* (2024) showed that WM can be maintained in the absence of persistent spiking [47]. Using single-trial analyses in nonhuman primates, they identified two coexisting maintenance- period states (an active spiking state and an inactive, low-firing state) and demonstrated that mnemonic information persisted through structured functional connectivity even during inactive phases. Their observation of stochastic transitions between states implies that trial averaging can create the illusion of continuous maintenance activity, aligning with our finding that heterogeneous single-neuron dynamics underlie apparent population-level persistence. It remains an open question whether the burst events we observe correspond to the active states described by Panichello *et al*. Here, by leveraging a cell-type classification focused on single-neuron bursting dynamics, we further extend this framework and show that transient bursts are predominantly driven by putative pyramidal neurons, suggesting a potential cellular substrate for these seemingly stochastic network transitions.

Our results further reveal a functional dissociation in how distinct cell types support WM, extending the population-burst analyses. Stable, content-specific bursting in putative pyramidal neurons aligns with their intrinsic burst-firing capabilities and provides a mechanistic basis for models of WM that rely on sparse, asynchronous coding [54, 55]. In contrast, dynamic, task- modulated bursting in putative interneurons underscores their roles in adaptive maintenance and circuit control [49, 52]. It remains to be determined specifically how these cell types interact to organize circuit-level dynamics of WM maintenance.

An important next step is to elucidate the mechanisms that support information retention during activity-silent or non-bursting periods. Prior work suggests that such latent states may arise from transient changes in functional connectivity, potentially mediated by short-term synaptic plasticity that modulates circuit excitability in the absence of sustained spiking [47, 76]. Future work should determine whether and how the interplay among diverse neuronal populations gives rise to activity-silent maintenance, providing a mechanistic bridge between transient bursting and the longer-lived network configurations that sustain WM.

In conclusion, our findings suggest that while persistent activity can emerge within distributed networks and remains a useful population-level signature of information maintenance, the nature of WM representations in local circuits is far more dynamic. We show that complementary interactions among heterogeneous neuronal populations support maintenance, with pyramidal neurons encoding information through sparse, burst-like events and interneurons modulating network dynamics to stabilize mnemonic representations.

## Methods

### Participants and data source

We analyzed the open-access dataset released by Kyzar *et al*. [60]. The cohort comprised 21 patients with intractable localization-related epilepsy who completed 41 recording sessions (age 17– 71 years; mean ± s.d., 36.10 ± 15.45). Participant recruitment, clinical procedures, and consent were conducted by the original investigators and approved by their institutional review boards, as reported in Kyzar *et al*. Data used here were de-identified and obtained from the DANDI Archive (https://doi.org/10.48324/dandi.000469/0.231213.2047).

### Psychophysical task and behavior

We used trials from the modified Sternberg working-memory (WM) task implemented by Kyzar et al. (Fig. 1a). Each trial comprised four epochs and began with fixation (900–1000 ms), followed by an encoding epoch in which participants memorized 1–3 images (load 1–3). For each participant, stimuli were drawn from a participant-specific set of five images defined in a prior screening session. Image sets differed across participants. Encoding was followed by a maintenance epoch of 2.5–2.8 s during which a white screen displaying “hold” appeared. A probe stimulus was then presented, and participants indicated whether it matched any encoded item by pressing a green (“yes”) or red (“no”) button. Participants were instructed to respond as quickly as possible, and the probe remained on screen until response. Reaction time (RT) was defined as the interval from probe onset to button press. Sixteen participants completed 135 trials and five completed 108 trials. Stimuli were presented in a pseudorandom order.

### Electrophysiology

Extracellular electrophysiological recordings were obtained using microwires embedded within hybrid depth electrodes (Ad-Tech Medical). Electrode targets were determined from on pre-operative structural MRI, and post-operative MRI and/or CT were co-registered to the MNI152-aligned CIT168 T1-weighted template for standardized localization. For visualization, recording sites were projected onto a 2D sagittal plane. Broadband potentials (0.1 Hz–9 kHz) were continuously sampled at 32 kHz using Neuralynx ATLAS or Cheetah systems. Channels exhibiting interictal epileptiform activity during a session were excluded. Spike sorting was performed with OSort after zero-phase-lag band-pass filtering (300–3,000 Hz). Spikes were detected using an energy-band method [60, 77, 78].

Across 21 participants, 902 single units were isolated from five regions: amygdala (AMY, *n* = 259), hippocampus (HPC, *n* = 190), dorsal anterior cingulate cortex (dACC, *n* = 171), presupplementary motor area (preSMA, *n* = 250), and ventromedial prefrontal cortex (vmPFC, *n* = 32; Fig. 1b). Spike times were provided as timestamp lists.

Spike times were segmented into task epochs: fixation, encoding (load 1–3), maintenance, and probe. For each neuron, firing rates were *z*-scored relative to the fixation period using the mean and s.d. across during fixation across trials for that neuron. For time-resolved analyses, peristimulus time histograms (PSTHs) used non-overlapping 50 ms bins (Fig. 1g), chosen to balance temporal resolution against trial-to-trial variability. A Gaussian filter was applied to the *z*-scored activity across bins (kernel *σ* = 1) to reduce high-frequency noise while preserving variability across task epochs, trials, and neurons.

### Identification of concept cells

Concept cells were defined as neurons whose firing during the first encoding epoch varied significantly with stimulus identity and was selective for a single identity [64, 79–81]. For each neuron recorded during the Sternberg WM task, spiking activity from the first encoding period across loads (1–3) was analyzed. Spike counts were computed in a 200–1000 ms window after stimulus onset [29]. To assess selectivity, we identified the two stimulus identities that elicited the highest mean firing rates (across trials in which that identity was presented in the first encoding period). Statistical significance of the difference in mean firing rate between these two top-ranked stimulus identities was tested using a nonparametric bootstrap test. For each neuron, trial-wise responses for the two identities were resampled with replacement (1,000 iterations), and on each iteration the mean-rate difference was computed to form a bootstrap distribution. The two-tailed *P* value was calculated as twice the smaller of the proportions of bootstrap differences above or below zero, testing whether the mean response difference between identities was statistically significant. Neurons with *P <* 0.05 were classified as concept cells. This nonparametric procedure accommodates unequal trial counts across stimulus identities and does not assume Gaussian response distributions within neurons.

### Generalized linear model (GLM) encoding analysis

To ensure consistent trial counts across participants and conditions, we constructed pseudo-populations for encoding analyses and population decoding by pooling neurons across participants while preserving within-participant labels [82–84]. Because trial numbers varied across participants, we matched to the minimum common count: 96 trials per participant matched on load for encoding analyses, and 75 trials per participant matched on presented stimulus identity for population decoding.

To quantify how individual neurons encode task-relevant variables, we fit a Poisson generalized linear model (GLM) to the spike counts of each neuron. For the maintenance epoch, we formed a trial-by-neuron array (pseudo-population) in which each entry was the spike count for neuron *n* on trial *t* separately within load 1 encoding and maintenance epochs. For each neuron, the trial-wise count vector served as the response in a Poisson GLM with canonical log link,

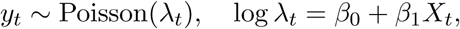

where *X_t_* was either memory load (1, 2, or 3) or RT (continuous).The predictor *X_t_* was *z*-scored prior to fitting for numerical stability. The coefficient *β*_1_ indexed each neuron’s sensitivity to the predictor.

Significance was assessed using a nonparametric permutation procedure (1,000 iterations). For each neuron, the trial-wise predictor *X_t_*was permuted *within* participants to preserve dependence structure, the GLM was refit for each permutation, and |*β*^∗^| was recorded to form a neuron-specific permutation-derived null distribution. A neuron was deemed significant if |*β*_1_| met or exceeded the 95^th^ percentile of its null (two-sided, *α* = 0.05). For group-level inference by region or by putative cell type (pyramidal, PY; interneuron, IN), the same permutations were used to generate a null distribution for the number of significant neurons. On each iteration and for each region or cell type, we counted the neurons exceeding their per-neuron cutoffs under scrambled labels and compared the observed count to this null using a right-tailed test. The *P* value was the fraction of null counts ≥ the observed [85].

### Population decoding analysis

To quantify stimulus-specific information in population activity, we trained a decoder on a pseudo-population (see above). For each participant, stimulus identities were labeled 1–5 (five-way classification). Pooling across participants provides a common feature space in which the decoder can learn weights for neurons that consistently signal, for example,“Image 1,” regardless of its visual identity across participants. For each trial (*T*) and neuron (*N*), spike times were binned into 250 ms windows with a 100 ms step size [86]. Spike counts were *z*-scored per neuron using fixation activity across all trials (as described in *Electrophysiology*). This yielded a *T* × *N* × (time bins) array. For decoding at a given time bin, we used the corresponding *T* × *N* slice as input features. Class labels were stored in a response vector *y* (five classes, one label per trial).

Linear support vector machines (SVMs) implemented via the Neural Decoding Toolbox [82] with LIBSVM [87]) were trained to predict the identity of the stimulus presented in the first encoding period, separately for each memory load. To assess temporal generalization, we performed cross-temporal decoding: for each training bin, the decoder was tested on every other bin, yielding a train×test time matrix of accuracies [88, 89]. Performance was evaluated with leave-one-out cross-validation (LOOCV) over trials. Chance level was 20% (five classes). For each train-test bin pair, we first computed a binomial test against the 20% chance level (two-sided, *α* = 0.05). We then controlled for multiple comparisons across the time-by-time grid using the Benjamini-Hochberg false discovery rate (FDR). To account for temporal dependence, we additionally generated an empirical cluster-duration null by repeating the full analysis 1,000 times with randomly permuted class labels in the target vector *y*. For each permutation, we applied the same binomial threshold, identified supra-threshold clusters, and recorded their temporal extent. Clusters in the empirical matrix whose duration did not exceed the 95^th^ percentile of this null distribution were removed. Surviving clusters were outlined in red in Fig. 2d, Fig. 6l, and Extended Data Fig. 1d.

We also trained and tested the SVM exclusively on maintenance-epoch activity to assess the reliability of memory representations across brain regions, participants, and loads. For cross-participant and cross-region analyses, we required at least three identified concept cells per participant to ensure stable estimates. To replicate prior work and examine bin-size effects [29], we first used 1,000 ms bins with a 250 ms step, then repeated the pseudo-population analysis while systematically varying bin width (250–1,000 ms; 25 ms step). Both analyses used LOOCV. Statistical significance of decoding accuracy was assessed via permutation: stimulus labels were randomly shuffled 1,000 times to obtain a null distribution of accuracies, and a *P* value was computed as the proportion of permuted accuracies exceeding the empirical value. FDR correction was applied within each family of comparisons.

### State-space analysis of population activity

We investigated neural population dynamics with demixed principal component analysis (dPCA) to compare encoding and maintenance during the WM task [29, 90, 91]. For dPCA, we constructed design matrices *X* from the neural activity during load 1 trials, separately for encoding and maintenance. Each matrix contained 30 trials, balanced across the five stimulus identities (6 per identity). For each trial, spike counts from all concept cells (*n* = 58; see *Identification of concept cells*) were assembled into a four-dimensional array with shape (trials × neurons × stimuli × time). Maintenance trials were truncated at 2.5 s for consistency across participants. Spikes were binned into non-overlapping 2 ms intervals and smoothed with a one-dimensional Gaussian kernel (*α* = 60; bin width = 120 ms). To account for baseline firing rate differences, each neuron’s activity was *z*-scored using its fixation-period mean and s.d. across trials.

To obtain robust, generalizable state-space estimates, we used stratified resampling. Within each iteration, trials were split into disjoint train and test sets such that each of the five stimulus identities contributed 3 of 6 trials to train and the remaining 3 to test (balance within identity). Splits were generated independently for each identity on each iteration, no trial appeared in both train and test sets, and each iteration used a unique split. This procedure was repeated 100 times, yielding 100 independent train/test combinations per epoch. Within each resampling iteration, we fit dPCA models to mean-centered, trial-averaged population activity from the *train* set, separately for encoding and maintenance, using “stimulus” and “time” as marginalization labels. Regularization parameters were selected by the package’s built-in cross-validation [90, 92, 93]. The corresponding *test* data were then projected into the low-dimensional dPCA space defined by the epoch-specific training model. In addition to projecting test data into their matched spaces (encoding → encoding; maintenance → maintenance), we also projected maintenance test data into the encoding-derived space to assess generalization of the encoding code to maintenance.

Stimulus separability was quantified for each condition and iteration by first computing the condition-averaged trajectory for each stimulus in the dPCA space, then computing the pairwise Euclidean distances between all stimulus centroids. These pairwise distances were summarized to yield a single separability score per iteration. To compare separability across conditions, we used a Kruskal–Wallis test followed by Dunn’s post hoc tests with FDR correction.

For analyses of temporal structure within maintenance, the same procedure was applied to the first (early) and second (late) halves of the maintenance period.

### Autocorrelogram-based characterization of single-neuron temporal dynamics

We characterized single-neuron temporal firing dynamics using spike-train autocorrelograms (ACGs). Spike times from all task epochs were aggregated for each of the 902 neurons and processed in MATLAB. For each neuron, we computed pairwise time differences between spikes as Δ*t* = *t_j_* − *t_i_*, for all *i* ≠ *j*, where *t_i_* and *t_j_* denote individual spike times. Self-coincidences (Δ*t* = 0) were excluded. These time differences were binned symmetrically around zero lag to produce a histogram of spike coincidences. We computed two ACGs per neuron to capture fine-scale and broader temporal structure:

- Narrow ACG: computed within a ± 50 ms window using 0.5 ms bins (201 bins total), capturing short-latency dynamics and burst-related peaks.
- Wide ACG: computed within a ± 500 ms window using 1 ms bins (1001 bins total), capturing rhythmic modulation such as theta-range oscillations.

Each ACG was divided by the total number of spikes and the bin width to yield a spike-density histogram. We then normalized each ACG by dividing all bin values by the maximum ACG amplitude for that neuron. This unit normalization rescales every ACG to [0, 1] to preserve temporal structure while enabling direct comparison of relative modulation and burst strength across neurons irrespective of absolute firing rates.

We fit each narrow ACG (using raw amplitudes, not unit-normalized) with bounded nonlinear least squares, using the default initialization in CellExplorer [73]. The resulting ACG-derived features (*τ*_rise_, *τ*_decay_, *R*^2^) were used in downstream analyses (e.g., cell-type classification and bursting analyses).

To ensure stable estimation, ACGs with poor fits (*R*^2^ *<* 0.3) were excluded from further analyses.

### Burst detection

We identified transient bursts of spiking activity using a procedure adapted from [59]. All analyses were performed within participant (i.e., no pseudo-populations). For each neuron, spike times were concatenated across all trials within a given memory load to obtain a continuous spike train for that load. Separate spike-time series were extracted for the encoding and maintenance periods, allowing burst activity to be analyzed independently in each task period. Burst detection depended on two parameters: the spike-count bin width and the Gaussian kernel width used to smooth the instantaneous rate. To constrain detection to biologically relevant timescales, the bin width was restricted to 50–150 ms, a range characteristic of burst dynamics [70]. Parameter selection was performed via a grid search over bin widths and Gaussian kernel widths using neural data from the encoding period only, comparing burst counts across parameter combinations. The combination Δ*t* = 70 ms (bin width) and *σ* = 40 ms (Gaussian kernel) maximized the burst count while remaining near the center of the search space, providing a balanced sensitivity-specificity trade-off (Extended Data Fig. 3a, [94]). These parameters were fixed for all participants to avoid participant-specific biases in detection sensitivity.

Within each participant, and for each memory load and epoch (encoding or maintenance), we binned and smoothed spike trains using the predetermined parameters. Let *x_t_* denote the smoothed bin value at time bin *t*. A bin *t* was defined as a local peak if:

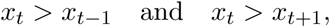

with a minimum inter-burst interval of 2 bins (140 ms). Peak prominence was computed as:

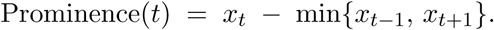

A local peak was classified as a burst if its smoothed firing rate exceeded the 90^th^ percentile of all time bins for that participant (thresholds computed separately within each epoch). In addition, we generated rate-matched Poisson surrogates for each neuron, matching its empirical firing rate on each trial. We simulated 100 surrogates per neuron and used the resulting null distribution of surrogate-derived burst counts for comparison with real data.

After within-participant burst detection, we repeated the procedure on neuronal groups to examine collective dynamics (e.g., concept cells and groups defined by ACG features). Neurons were median-split within each participant based on mean ACG amplitude and the decay time constant (*τ*_decay_). This yielded four equal-sized groups per participant: Highest ACG, Longest *τ*_decay_, Lowest ACG, and Shortest *τ*_decay_. To ensure stable estimates and comparability across participants, we excluded participants contributing fewer than 5 neurons to a given group [95]. For each participant and neuronal group, we computed (i) the total number of bursts in the real data and (ii) the mean number of bursts across that matched set of 100 Poisson surrogates. We summarized these participant-level values across the cohort by reporting the mean and s.e.m. for both real and surrogate burst counts. To test whether real-data burst counts exceeded chance, we compared participant-level real versus surrogate counts with paired Wilcoxon singed-rank tests. Multiple comparisons were controlled with FDR correction, with *α* = 0.05.

We then examined how bursting evolves across task epochs and memory loads. Using the neuronal groups defined above, we performed the analysis separately for loads (1–3). To capture temporal heterogeneity within maintenance, we divided this epoch into three consecutive 1 s win- dows (M1, M2, M3) and computed burst counts within each window. For each load, we thus obtained participant-level burst-count distributions across encoding, maintenance (M1–M3), and probe. Distributions were visualized with box plots and compared using Wilcoxon signed-rank tests (within participant and neuronal group), with significance thresholds adjusted for multiple comparisons (FDR).

### Putative cell-type classification

To characterize neurons as putative pyramidal neurons (PY) or interneurons (IN), we used autocor- relogram (ACG)–derived metrics and a data-driven clustering approach. While such classification is common in rodent studies, its application to human data has been limited by the noise and sparsity of single-neuron recordings and the lack of established ground-truth labels [58, 96]. We view this step as essential for more accurately estimating the the neural dynamics underlying memory main- tenance. Guided by prior work, including a recent human study [57], and by our initial observation of prominent burst dynamics in a subset of neurons, we focused the classification on the burstiness dimension of cell-type differences. This dimension is directly observable in the ACG and provides a clear, interpretable axis for classification. Specifically, we used three features with robust cell-type differences reported in prior work: firing rate, mean normalized ACG amplitude, and the ACG rise time constant (*τ*_rise_) [57, 71, 72]. The latter two features capture the burstiness and single-neuron temporal dynamics, which are relevant to computational roles in memory maintenance. Because reference values for these metrics are not established in humans, fixed manual thresholds were not used. To accommodate the noisier conditions of our recordings, we adopted spectral clustering rather than the Gaussian Mixture Model (GMM) approach used previously [57], as spectral clus- tering is more robust to noise and makes fewer parametric assumptions about cluster shape. We also used the normalized mean ACG to emphasize the temporal shape [97]. The resulting clusters were interpreted as PY and IN.

To further quantify differences between clusters, we compared bursting-related metrics. Specif- ically, we computed the CellExplorer *burstiness index* as defined by Royer *et al*. [98] and the local coefficient of variation, CV_2_, for each neuron [99].

Bursting analysis was performed as defined in *Burst detection*, with one modification: we ex- cluded participants contributing fewer than 3 neurons to a given neuronal group, given the small pyramidal sample size.

Because our human recordings lack ground-truth cell-type labels, we evaluated the approach on a CellExplorer reference dataset with ground truth, comprising 184 parvalbumin (PV) and 115 somatostatin (SST) interneurons, alongside 44 pyramidal neurons [73]. Using *τ*_rise_, normalized mean ACG amplitude, and firing rate as inputs to spectral clustering, 333 of 343 neurons were correctly assigned, with only one pyramidal neuron and nine interneurons (PV or SST) misclassified (Extended Data Fig. 4a). These results support the accuracy and cross-dataset generalizability of the method.

### Statistical analyses

Descriptive statistics are reported as mean ± s.d., except reaction times (RT), which are reported as median with inter-quartile ranges (IQR). Unless otherwise stated, bar plots show mean ± s.e.m., and box plots show the median and IQR. All main-figure analyses, except for the Poisson GLM analysis of encoding, focus on load 1 trials. Results for load 2 and 3 are provided in the *Supplementary Information*.

All statistical analyses were performed with custom code in Python (and in MATLAB for cell- type classification; see above). Significance was assessed with nonparametric permutation tests. Specifically, labels were randomly shuffled to generate a null distribution of the test statistics (e.g., decoding accuracy, difference in means), and *P* values were computed as the proportion of permutations exceeding the observed value. The number of shuffles was 1,000 unless otherwise specified (analyses using 100 shuffles are noted in the relevant subsections).

For pairwise comparisons, we used the Wilcoxon signed-rank test. Multiple comparisons were controlled using the Benjamini-Hochberg false-discovery-rate (FDR) procedure with a corrected significance threshold of *P <* 0.05 [100]. All tests were two-sided unless stated otherwise. Tests of decoding against chance were one-sided: cross-temporal decoding used a binomial with cluster- level thresholding [101], and maintenance-only decoding used a permutation test with 1,000 label shuffles.

---

**Extended Data Fig. 1.**
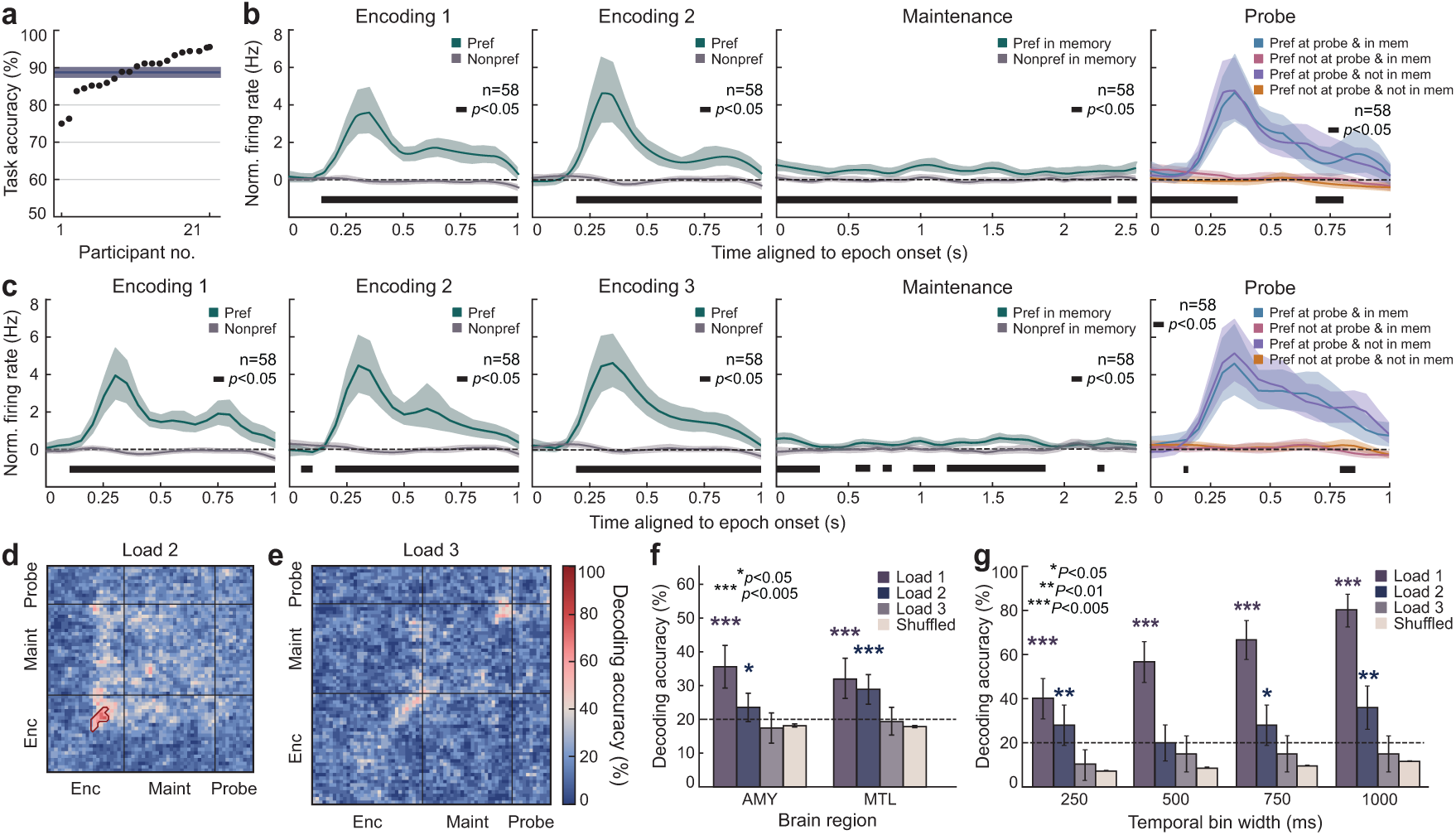
**| Stimulus decoding across task epochs at higher load (loads 2–3). a,** Response accuracy by patient, rank-ordered from lowest to highest. The solid line shows the group mean and the shaded band shows s.d. across patients. **b,** Mean normalized firing of all concept cells (*n* = 58) across task epochs for load 2 trials when the preferred image was presented versus when the nonpreferred image was presented during the encoding epoch. Shaded bands indicate 95% confidence intervals. Black bars indicate time bins with significant differences between conditions (permutation test, *P <* 0.05). When averaged, concept-cell activity appears persistent during maintenance. **c,** As in **b,** for load 3 trials. Persistent activity was not observed during the maintenance epoch. **d,** Cross-temporal decoding for load 2 trials. The y-axis denotes training time bins; the x-axis denotes testing time bins. Color indicates accuracy for identifying the encoded stimulus. Red contours mark clusters with accuracy above chance (permutation test, *P <* 0.05). Chance was set to [1/number of stimulus classes] (20%). Cluster sizes expected under the null were estimated from 1,000 label-shuffled iterations and subthreshold clusters were omitted. **e,** As in **d,** for load 3 trials. **f,** Decoding of stimulus identity during the maintenance epoch from concept cells in AMY and MTL, pooled across participants, for load 2 trials. Shuffled-label controls are shown for comparison. The dashed line marks chance (20%). Accuracy exceeded chance in AMY and in MTL (permutation test, *P <* 0.005). **g,** Decoding accuracy for stimulus identity during maintenance as a function of the temporal bin width. A linear SVM with LOOCV was applied to load 1, load 2, and load 3 trials across bid widths. Shuffled-label controls are shown for comparison. The dashed line marks chance (20%). Asterisks indicate accuracies above chance or differences between conditions (permutation tests). \**P <* 0.05; \*\**P <* 0.01; \*\*\**P <* 0.005.

**Extended Data Fig. 2.**
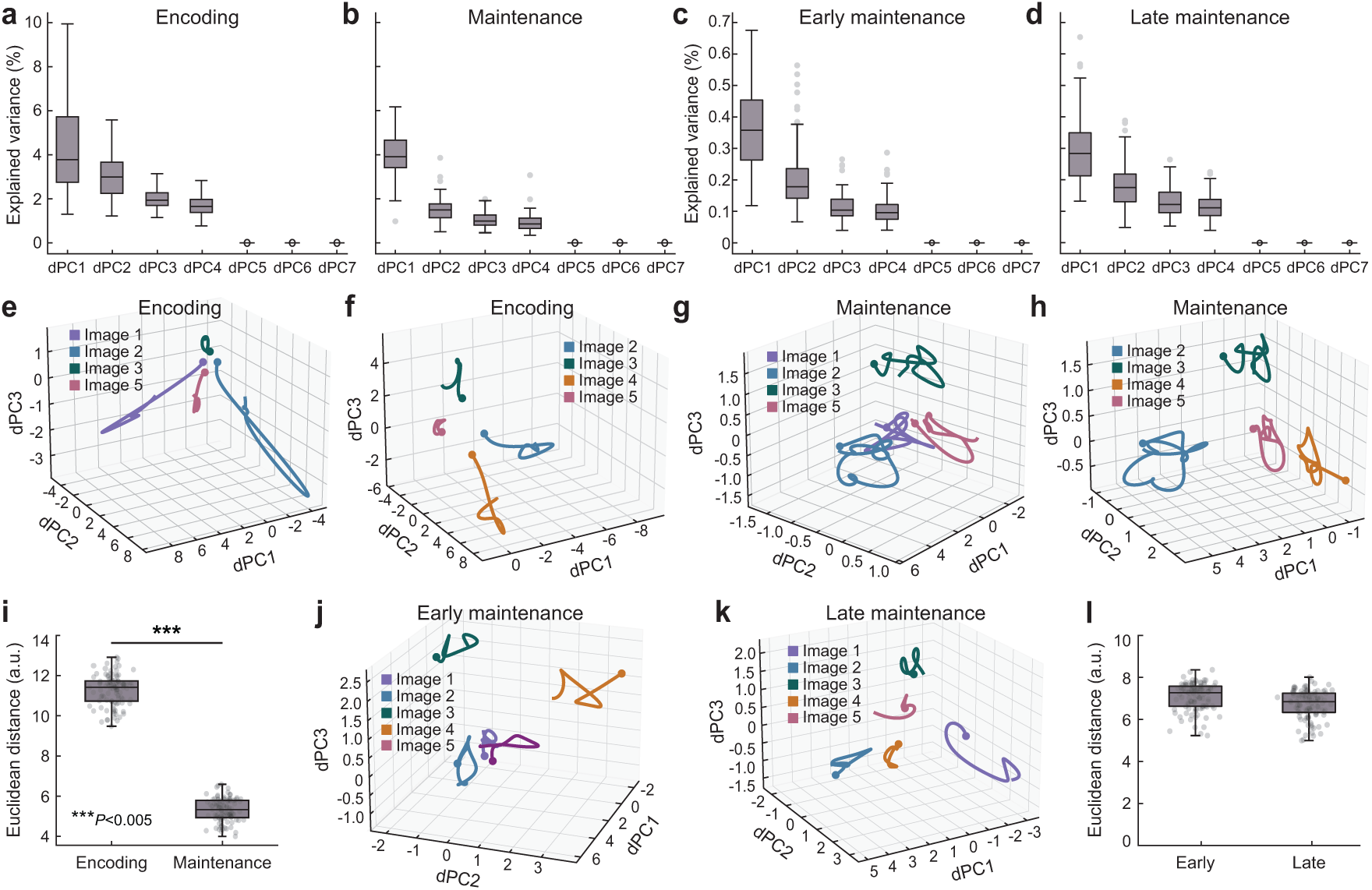
**| Encoding and maintenance occupy distinct neural subspaces. a,** Explained variance of demixed principal components (dPCs) for models fit on data from the encoding epoch. Box plots show the median across 100 training shuffles. Bottom and top edges indicate the lower and upper quartiles. Whiskers extend to 1.5 × interquartile range. **b,** As in **a,** for the maintenance epoch. In **a**–**b**, the first four dPCs explain most of the variance. **c,** Explained variance of dPCs fit on data from the first half of maintenance epoch. Box plots show the median across 100 training shuffles. Bottom and top edges indicate the lower and upper quartiles. Whiskers extend to 1.5 × interquartile range. **d,** As in **c,** for the second half of the maintenance epoch. In **c**–**d**, the first four dPCs capture most of the variance. **e–f,** dPCA fit to encoding training data separates activity for the five stimulus identities (first three dPCs shown. Trajectories are split across two panels for clarity. **g–h,** Neural trajectories from maintenance train trials, projected into the maintenance state space, show clear separation by stimulus identity. Trajectories are split across two panels for clarity. **i,** Pairwise Euclidean distances between averaged population responses for each stimulus identity, computed from training trials for encoding in the encoding state space and for maintenance in the maintenance state space, show greater stimulus separation during encoding than during maintenance (Kruskal-Wallis with Dunn’s post hoc, *P* = 2.75 × 10*^−^*^22^; see *Methods*). **j,** dPCA fit on data from the first half of the maintenance epoch. **k,** As in **j,** for the second half of the maintenance epoch. In **j**–**k**, the state spaces separate activity for the five stimulus identities. **l.** Pairwise Euclidean distances between averaged population responses for each stimulus identity, computed from training trials for early maintenance in the early-maintenance state space and for late maintenance in the late-maintenance state space, indicate similar stimulus separation (Kruskal-Wallis with Dunn’s post hoc, *P >* 0.05). \*\*\**P <* 0.005.

**Extended Data Fig. 3.**
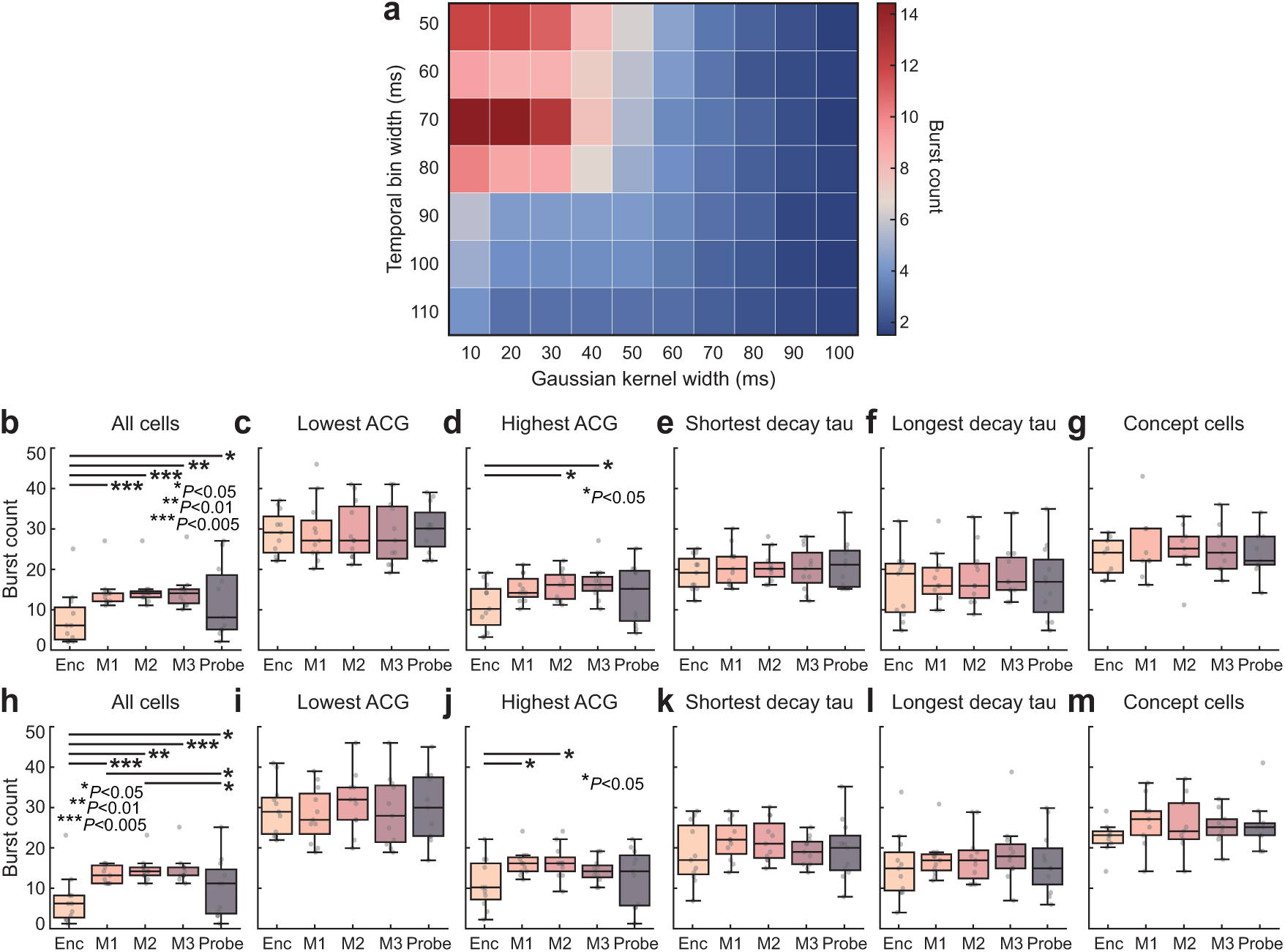
**| Bursting dynamics are consistent at higher WM loads. a,** Grid search over Gaussian kernel width (x-axis) and temporal bin width (y-axis). Parameters were selected to balance noise reduction and temporal resolution, yielding a chosen parameter setting in the central region of the grid. *Panels **b-g.*** Burst rates by neuronal groups for load 2 trials, across the second encoding period, maintenance, and probe. Wilcoxon signed-rank tests were used, with multiple comparisons controlled. Neuronal group definitions, burst detection, and rate normalization and detailed in *Methods*. **b,** All cells combined: burst rate during maintenance is higher than during encoding, consistent with a task-dependent temporal reorganization of spiking (*P*_max_ = 0.020). **c,** Low ACG cells: no significant change in bursting from encoding to maintenance (*P >* 0.05), indicating that low ACG neurons do not drive the increase seen at the population level. **d,** High ACG cells: maintenance shows a robust increase in bursting relative to encoding (*P*_max_ = 0.020), suggesting that high ACG neurons contribute disproportionately to the population-level effect. **e,** Cells with a short *τ*_decay_: no significant difference in bursting between encoding and maintenance (*P >* 0.05). **f,** Cells with a long *τ*_decay_: no significant difference in bursting between encoding and maintenance (*P >* 0.05). **g,** Concept cells: no significant change in bursting between encoding and maintenance (*P >* 0.05). **h–m,** As in **b–g,** for load 3 trials. Box plots: center lines, median; bottom and top edges, lower and upper quartiles; whiskers, 1.5 × interquartile range. \**P <* 0.05, \*\**P <* 0.01, \*\*\**P <* 0.005.

**Extended Data Fig. 4.**
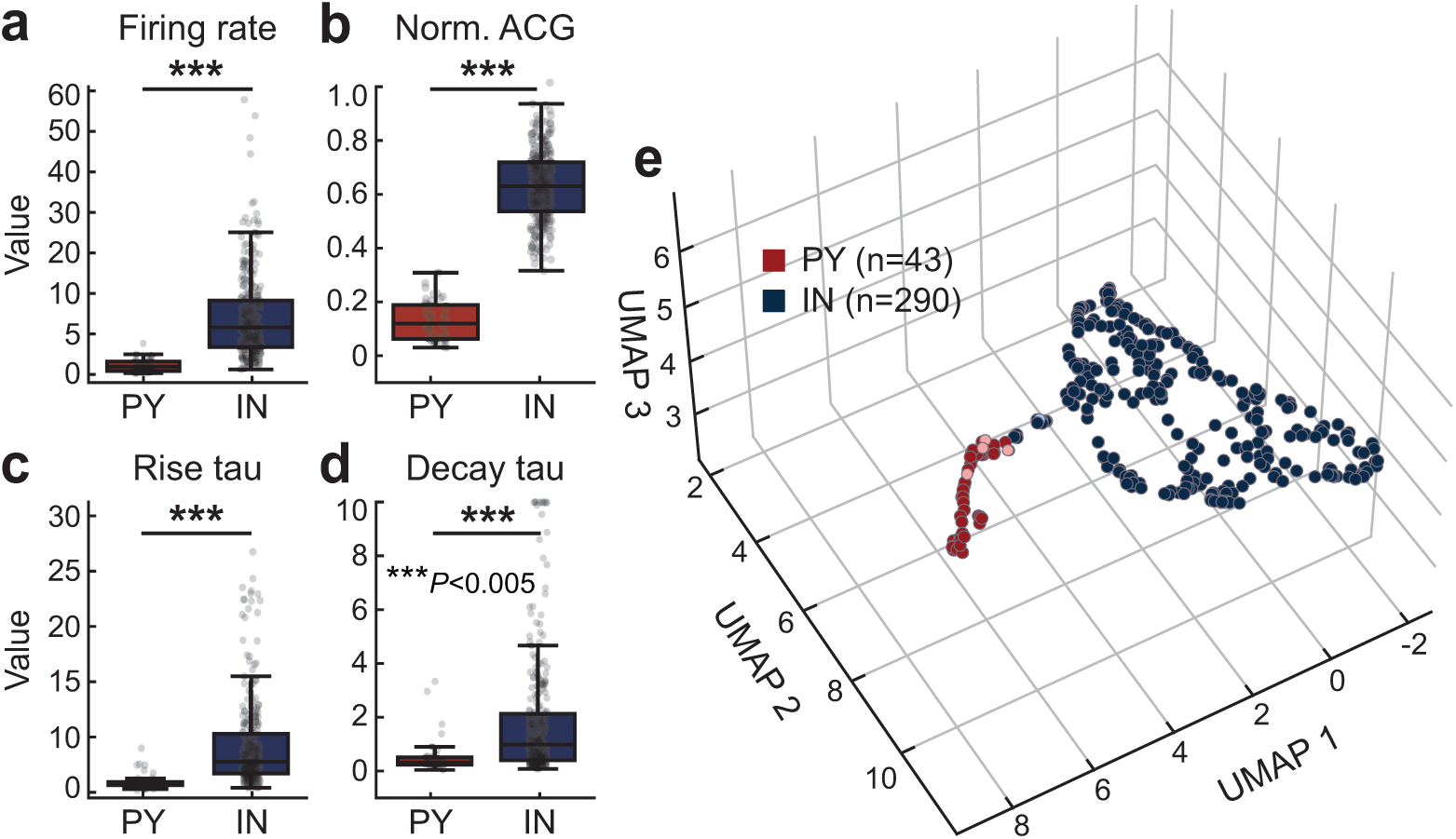
**| Validation of the cell-type classifier on a dataset with ground-truth labels. a,** Firing rate by putative cell type. Interneurons had higher rates than pyramidal neurons (Wilcoxon rank-sum; *P* = 2.49 × 10*^−^*^22^). **b,** Mean normalized ACG by putative cell type. Interneurons had higher mean ACG than pyramidal neurons (Wilcoxon rank-sum, *P* = 1.44 × 10*^−^*^25^). **c,** ACG rise time constant (*τ*_rise_) by putative cell type. Interneurons had higher *τ*_rise_ than pyramidal neurons (Wilcoxon rank-sum, *P* = 1.68 × 10*^−^*^17^). **d,** ACG decay time constant (*τ*_decay_) by putative cell type. Interneurons had higher *τ*_decay_ than pyramidal neurons (Wilcoxon rank-sum, *P* = 1.10×10*^−^*^9^). **e,** Spectral clustering using firing rate, mean ACG, and ACG rise time constant (*τ*_rise_), identified two groups in a labeled dataset [73]. A UMAP projection for visualization shows points colored by the spectral clustering assignments, corresponding to putative pyramidal (PY, dark red) and interneuron (IN, dark blue) populations (PY: *n* = 43; IN: *n* = 290). Pink and light blue points mark misclassified neurons (PY: *n* = 1; IN: *n* = 9). Overall, 97% of labeled cells were classified correctly (see *Methods*). Box plots: center lines, median; bottom and top edges, lower and upper quartiles; whiskers, 1.5 × interquartile range. \*\*\**P <* 0.005.

**Extended Data Fig. 5.**
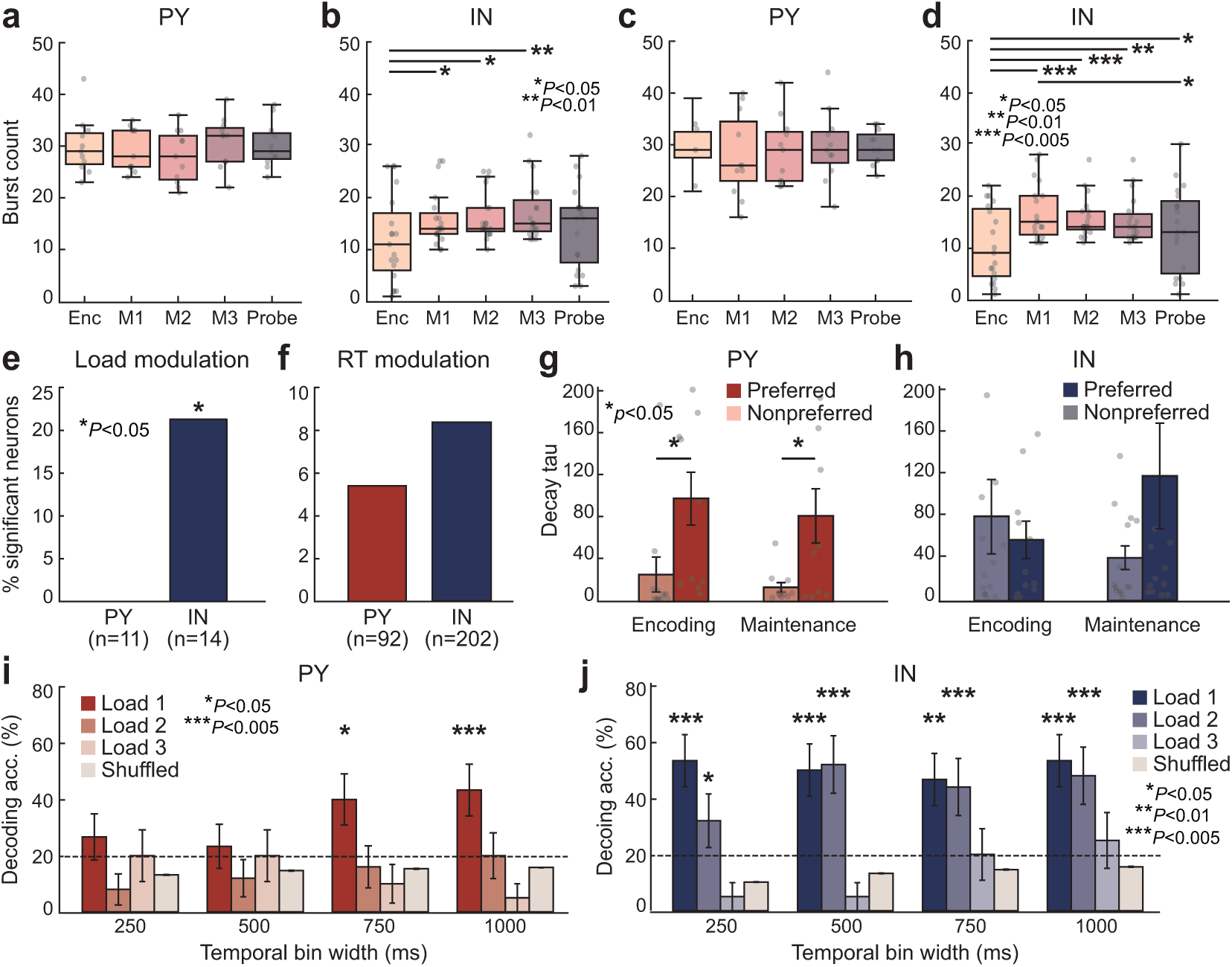
**| Differential dynamics of pyramidal neurons and interneurons at higher loads. a,** Burst counts across task epochs (encoding; maintenance M1, M2, M3; probe) in load 2 trials for pyramidal neurons (PY), where M1, M2, and M3 denote the first, second, and final thirds of the maintenance epoch. **b,** Burst counts across epochs for interneurons (IN). IN exhibit epoch-dependent changes (Wilcoxon signed-rank, *P*_max_ = 0.033). **c,** As in **a,** for load 3 trials. **d,** As in **b,** for load 3 trials. **e,** Proportion of concept-selective PY and IN with load-modulated firing during maintenance (permutation test, *P*_PY_ *>* 0.05, *P*_IN_ = 0.033). **f,** Proportion of PY and IN with reaction time (RT)-modulated firing during maintenance (permutation test, *P >* 0.05). **g,** ACG Decay time constant (*τ*_decay_) on preferred and nonpreferred trials across encoding and maintenance for concept-selective PY. **h,** As in **g,** for concept-selective IN. Concept-selective PY, but not IN, show a change in (*τ*_decay_) on preferred trials (Wilcoxon signed-rank, uncorrected *P*_max_ = 0.029). **i,** Decoding of stimulus identity during maintenance from PY concept cells for load 1, 2, and 3 trials as a function of temporal bin width (250, 500, 750, and 1,000 ms). Bin width strongly affects PY accuracy in load 1 trials. Accuracy does not exceed chance level for load 2 or load 3 trials (permutation tests, all *P >* 0.05). **j,** Decoding accuracy during maintenance from IN concept cells for load 1, 2, and 3 trials as a function of temporal bin width. IN decoding accuracy remains above chance across all bin widths in both load 1 and load 2 trials, and is less sensitive to bin width than PY (permutation tests, *P*_max_ = 0.015). *Panels **i–j***. Shuffled-label controls are shown for comparison. The dashed line denotes chance (20%). Asterisks indicate accuracies above chance. \**P <* 0.05, \*\**P <* 0.01, \*\*\**P <* 0.005.

